# Formation and recycling of an active epigenetic mark mediated by cell cycle-specific RNAs

**DOI:** 10.1101/2021.10.27.466094

**Authors:** Alexander K. Ebralidze, Simone Ummarino, Mahmoud A. Bassal, Haoran Zhang, Bogdan Budnik, Emanuele Monteleone, Dennis Kappei, Yanjing V. Liu, Danielle E. Tenen, Rory Coffey, Mee Rie Sheen, Yanzhou Zhang, Anaïs Wanet, Bon Q. Trinh, Valeria Poli, Vladimir Espinosa Angarica, Roberto Tirado Magallanes, Touati Benoukraf, Colyn Crane-Robinson, Annalisa Di Ruscio, Daniel G. Tenen

## Abstract

The mechanisms by which epigenetic modifications are established in gene regulatory regions of active genes remain poorly understood. The data presented show that the establishment and recycling of a major epigenetic mark, the acetylated form of the replacement histone H2A.Z, is regulated by cell cycle-specific long noncoding RNAs encoded in regions adjacent to the promoters of active genes. These transcripts, termed *SPEARs* (S Phase EArly RNAs), are induced in early S phase: their expression precedes that of the downstream genes on which they exert their regulatory action. *SPEARs* drive the modification and deposition of the acetylated form of histone H2A.Z by bringing together the replacement histone and the histone acetyl transferase TIP60. This widespread bimodal pathway constitutes a novel RNA-mediated mechanism for the establishment of epigenetic marks and cell-specific epigenetic profiles, thereby providing a unifying explanation for the accuracy and persistence of epigenetic marks on chromatin.

---

A wealth of epigenomics data have recently provided meaningful biological and clinical information but little attention has been paid to the mechanisms for the establishment (‘writing’) and the maintenance of specific epigenetic patterns. Epigenetic aberrations are a frequent feature of deleterious conditions such as cancers and other genetic diseases^1^. In particular, subsets of histone variants and their post-translational modifications have emerged as prominent players among other epigenetic marks. The histone H2A.Z (the major H2A variant), for instance, is associated with gene activation^2–5^ by means of the balance between the acetylated *vs*. the non-acetylated forms at the promoters of genes^6–8^. Acetylation of H2A.Z is catalyzed by the histone acetyltransferase TIP-60 subsequent to its inclusion within nucleosomes^9^ but it remains unexplained how the balance between the acetylated and unmodified forms is inherited and preserved.

The concepts underlying epigenetic inheritance have recently undergone significant change. A number of studies demonstrated that epigenetic marks can cross generation borderlines^10–12^ but the mechanistic aspects of this intergenerational transmission of information, the so-called ‘epigenetic memory’, remained unclear. In particular, how is the balance between the acetylated and unmodified forms inherited and preserved. Here, we show that cell type-specific RNAs are conduits of the accurate recycling of the epigenetic mark and thereby provide transgenerational design marks for regulation of the transcriptional activity of the respective genomic loci.

Widespread transcriptional activity across the mammalian genome results in the production of many functional noncoding RNAs (ncRNAs)^13–18^ that play critical roles in multiple biological processes^18–29^. Such RNAs are essential components of the chromatin architecture^30–32^ that is shaped by epigenetic marks such as DNA methylation, histone modifications, nucleosome positioning, and the incorporation of histone variants into nucleosomes. Despite their well-documented role, few of these RNAs have been fully characterized and the majority still lack a functional classification.

The overall effect of these ncRNAs is to regulate cell-type specific gene expression at both the transcriptional and post-transcriptional level, including: (i) chromatin remodeling by recruiting complexes that lead to epigenetic changes (*e.g*., polycomb repressive complex (PRC2)–mediated transcriptional regulation^33^; (ii) transcriptional interference by directly interacting with ubiquitous or tissue specific transcription factors (*e.g*., dissociation of the pre-initiation complex through sequestration of the general transcription factor IIB^34^ or Sp1 in Myotonic Dystrophy cases^24^; or (iii) splicing interference by affecting the distribution of serine/arginine splicing factors^35^.

A targeted and global genomic approach is used here to demonstrate that the deposition and modification of the active *versus* inactive H2A.Z mark is mediated by locally induced cell-cycle specific RNAs, termed *SPEARs* (**S P**has**e EA**rly **R**NAs).

## RESULTS

### Expression of newly minted promoter RNAs correlates with that of neighboring genes

Since the majority of active genes undergo replication in early S phase, which is followed by formation of the appropriate active chromatin^36^, we reasoned that transcripts generated in early S phase might control expression of the respective mRNAs by playing a role in reassembly of the chromatin. Furthermore, we hypothesized that the epigenetic balance between acetylated and unmodified forms of H2A.Z could be maintained by locally induced ncRNAs.

To establish the global occurrence of such newly minted RNAs, nascent S-phase transcripts were captured and sequenced (nasRNA-Seq). Synchronized human HL-60 cells were labeled with the ribonucleotide homolog 5-ethynyl uridine (EU) for one hour upon release into S phase. The collected RNAs were then biotinylated by click chemistry, isolated on streptavidin beads and deep-sequenced to produce nasRNA-Seq libraries^37,38^ (Extended Data Fig. 1a and Methods for details). Analysis of such libraries demonstrated a strong correlation of expression levels between nascent transcripts arising from gene promoter regions (“nascent promoter RNAs”) and those of their linked genes (Fig. 1a). In particular, such correlations are more prominent for nascent transcripts arising from the promoter regions of highly expressed genes (Fig. 1a). Among the early expressing genes marked by the presence of early S-phase promoter RNAs we identified *c-MYC* (Extended Data Fig. 1b,c), the oncogene most frequently altered in cancers^39^, myeloid master regulator transcription factor *PU.1* (ref. ^40^) and proto-oncogene *MYB* (ref. ^41^) (uninterrupted sequence segments are in Supplementary Data #2); a list of all gene loci expressing early promoter RNAs is in Supplementary Data #1. The *c-MYC* early S-phase promoter transcripts were shown to be represented by ~13 copies in the nucleus of HL-60 cells (Extended Data Fig. 1d) and were mapped by primer extension and 5’, 3’-RACE (Extended Data Fig. 1e,f).

**Fig. 1.**
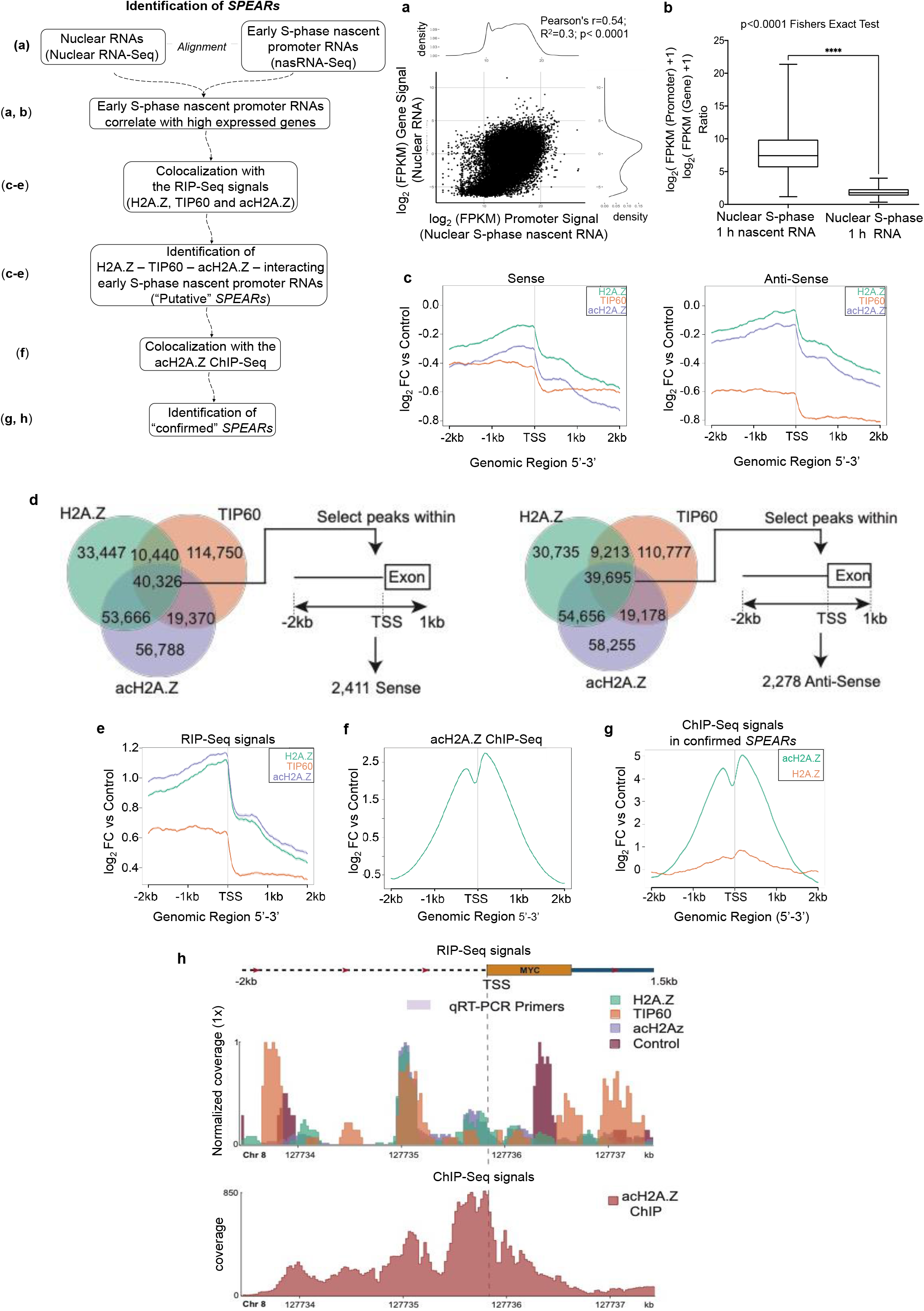
Characterization of the promoter RNAs induced in the first hour of S phase. **a**, Genome-wide analysis of the nascent nuclear RNAs (nasRNAs). Correlation between the nascent promoter RNA level and the expression of their respective coding genes. Pearson’s correlation score is 0.54, with an R^2^ value of 0.3, and p-value < 0.0001. **b**, Ratio of Promoter to Exon expression of S-phase nascent nuclear transcripts and of S-phase total nuclear transcripts, both in the first hour. P-value < 0.0001. **c**-**h**, ***SPEARs* interactions with H2A.Z, acH2A.Z and TIP60. c**, Read coverage plots for sense (forward) and antisense (reverse) strands for the region −2 kb to +2 kb surrounding transcriptional start sites (TSS). Note enhanced signal in upstream 2 kb region compared to downstream 2 kb region for all transcripts on both strands. **d**, Left Panel (Sense strand) a total of 40,326 RIP peaks overlapped the H2A.Z (green), TIP60 (orange) and acH2A.Z (purple) RIPs using a merge window of 1kb. From the 40,326 overlapping peaks those having a center located in the range −2 kb -TSS- +1 kb were selected, resulting in 2,411 being retained. Right Panel (Anti-Sense strand) a total of 39,695 RIP peaks overlapped the H2A.Z (green), TIP60 (orange) and acH2A.Z (purple) RIPs using a merge window of 1kb. From the 39,695 overlapping peaks those having a center located in the range −2 kb -TSS- +1 kb were selected, resulting in 2,278 being retained. Altogether, 4,689 promoter RNAs were identified from a total of 4,351 genes across both strands. **e**, RIP-Seq signals: read coverage plot for the region −2 kb to +2 kb surrounding TSSs. A significant enrichment of transcripts is seen in the upstream 2 kb region. **f**, Total acH2A.Z ChIP-Seq signals in the same region as RIP-Seq signals in Panel E, highlighting the increase in promoter signal at the same position as the S phase nascent promoter RNAs. **g**, H2A.Z and acH2A.Z ChIP-Seq signals overlapped just with confirmed *SPEARs*. **h**, Upper Panel - A normalized genomic coverage plot showing identified *SPEARs* in the region of the *c-MYC* TSS. The normalized coverage signals for each sample are overlaid to show good co-mapping of signals for H2A.Z (green), TIP60 (orange) and acH2A.Z (purple). The results of the qRT PCR (using the indicated amplicon) are shown in Figure S2CD. Bottom Panel: the acH2A.Z ChIP-Seq signals in the same region of the *c-MYC* gene, highlighting increased signal at the same positions as the *c-MYC SPEARs*.

### A fraction of early S-phase promoter RNAs interacts with H2A.Z, acH2A.Z and TIP60

Given the close proximity of promoter RNAs to TSS and the growing body of literature showing that highly expressed genes are marked by the acetylated form of the variant histone H2A.Z (acH2A.Z) (ref. ^5^), the possibilities of a link between early S-phase promoter RNAs and H2A.Z/acH2A.Z was investigated by: (i) Ribonucleoprotein (RNP) pull-down experiments followed by mass spectrometry; and (ii) RNA immunoprecipitations followed by RNA sequencing (RIP-Seq).

To design the RNA probes for RNP pull-down, the sequences of three early S-phase promoter RNAs, identified by nasRNA-Seq (Supplementary Data #1) and derived from gene loci *c-MYC, PU.1*, and *MYB*, were verified by primer extension and 5’, 3’ RACE (Figures S1E and S1F). The approximately 500 nt uninterrupted sequence segments (Supplementary Data #2) were cloned under a T7 RNA polymerase promoter to enable expression of biotinylated sense and antisense early promoter RNAs probes for RNP pull-down experiments (Extended Data Fig. 2a). Recovered RNPs interacting with both the antisense and sense early S-phase promoter RNAs probes and a negative control (D-Biotin) were analyzed by mass spectrometry. Among peptides pulled down by the early promoter RNAs probes were several corresponding to H2A.FZ/ H2A.FV (Supplementary Data #3). It was not possible to distinguish the very homologous forms of H2A.FZ *vs*. H2A.FV, and no peptides containing acetylation sites were detected, due to the relatively low fragment coverage. Other peptides pulled down by the early promoter RNA probes corresponded to histones H2A, H2B, H3, H4 and H1. No peptides corresponding to H2A/H2A.Z or other histones were detected by the negative control probe (Supplementary Data #3).

It is possible that early S-phase promoter RNAs exert their function through direct interaction with H2A.Z and the histone acetyl-transferase (HAT) TIP60, the enzyme reported to acetylate H2A.Z^9,42^. To directly test the interaction of early S-phase promoter RNAs with H2A.Z, acH2A.Z, and TIP60, RIP-Seq experiments were performed using antibodies to H2A.Z (i.e. recognizing all forms of the histone), to acetylated H2A.Z (acH2A.Z), and to TIP60 (Extended Data Fig. 2b). A significant overlap of RIP-Seq peaks was observed from all three immunoprecipitations and considerable co-localization of the three data sets was detected in the regions around the mRNA TSSs. In particular, a significant enrichment was observed of both sense and antisense transcripts in the 2 kb window upstream of TSSs (total number of early S-phase nascent promoter RNAs: 4689; 2,411 and 2,278 transcripts on the sense and antisense strands, respectively; Fig. 1c,d). At this stage, all early promoter RNAs which did not have detectable expression above background in the nasRNA-Seq library were discarded, which restricted the list of early S-phase nascent promoter RNAs to 3,560 (1,810 and 1,750 transcripts on the sense and antisense strands, respectively) (Fig. 1e). To assess the functional impact of the interaction of the early S-phase nascent promoter RNAs with TIP60, H2A.Z, and acH2A.Z, the identified 3560 early S-phase nascent promoter RNAs were aligned with the acH2A.Z ChIP-Seq library. To ensure consistency and stringency, only gene loci with a detectable acH2A.Z ChIP signal in the 2 kb upstream window of the TSS were accepted (Fig. 1f). This identified 2363 promoter RNAs (1209 sense and 1154 antisense transcripts) which are expressed in the first hour of S phase and which overlap with the signals of all four data sets: the three RIP-Seq libraries (antibodies to H2A.Z, TIP60, and acH2A.Z) and the acH2A.Z ChIP-Seq library. The co-location of acH2A.Z ChIP-Seq signals in the same genomic region as the signals from the three independent RIP-Seq experiments suggests a functional role of the early S-phase nascent promoter RNAs in positioning the acetylated H2A.Z marks. We termed these 2,363 early S-phase nascent promoter RNAs *SPEARs* (**S P**has**e EA**rly **R**NAs). Comparison of the H2A.Z and acH2A.Z ChIP-Seq data sets implies that it is the acetylated form of H2A.Z that is largely bound to the *SPEARs* (Fig. 1g). Taken together, the histone immuno-precipitations and the RIP experiments indicate: i) a triple interaction of *SPEARs* with H2A.Z, acH2A.Z, and with TIP60 in the 1 kb upstream of the TSS (Extended Data Fig. 2c); and ii) a correlation of *SPEARs* expression level with that of the linked genes.

In summary, identification of 2,363 *SPEARs* linked to expressed genes, direct correlation of *SPEARs* expression levels with those of the corresponding genes, and, importantly, with the occupancy of acH2A.Z within the respective *loci* (Fig. 1f-h), implies a global involvement of *SPEARs* in cooperation with TIP60 and H2A.Z/acH2A.Z in establishing an active expression mode at the corresponding genes. This warranted further examination.

### *SPEARs* carry common binding motifs

To further test the *SPEARs*-H2A.Z/TIP60 relationship, we performed motif discovery analysis on the *SPEARs* (see Methods for details). Among the twenty-five identified motif candidates, three motifs (##3, 5, and 9; Supplementary Data #4) exhibited the most significant enrichment among the *SPEARs*-regulated gene loci. Importantly, no similar motifs were found within the largest classes of nuclear RNAs, i.e. transcripts arising from SINEs and LINEs, nor from ribosomal RNA genes (see Methods for details). Among the top three candidates, motif 9 (corresponding to the RNA oligonucleotide RM9A; Fig. 2a and Extended Data Fig. 3a) was ranked as the strongest motif enriched in the *c-MYC SPEARs* sequence. This motif was enriched in acH2A.Z and TIP60 RIP-Seq overlapping peaks (13%), located near the peak centers (Fig. 2b).

**Fig. 2.**
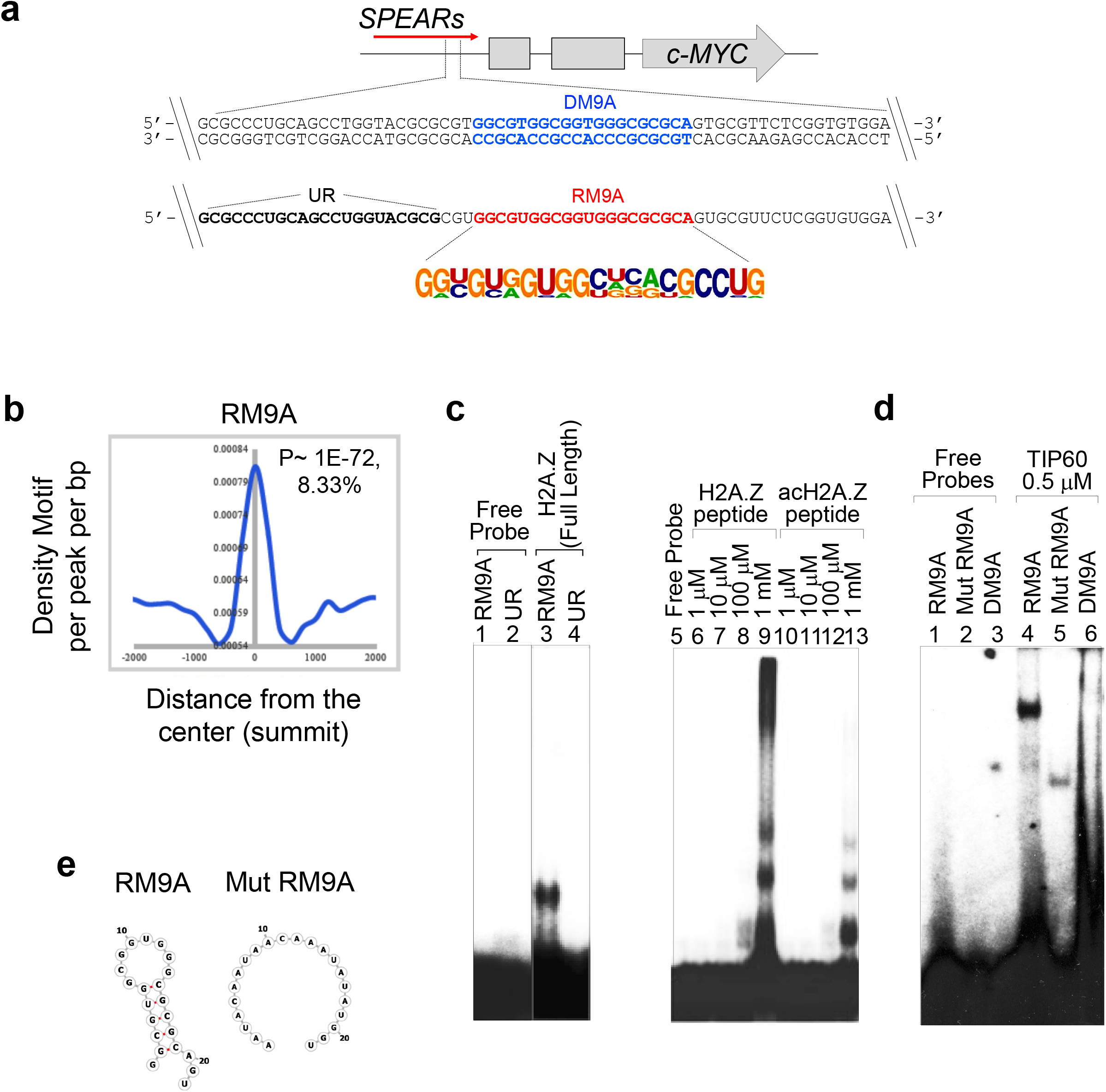
Identification of common binding motifs in *SPEARs*. **a**, The sequence of the common binding motif “9” (RNA oligonucleotide RM9A) and the unrelated RNA oligonucleotides (UR) within the *c-MYC SPEARs*, along with the corresponding motif “9” DNA duplexes (DM9A) are indicated. **b**, Distribution of the RM9A motif in acH2A.Z and TIP60 RIP-Seq overlapping peak regions. Significance and percent enrichment indicated. **c**, Left panel: REMSA gel for *c-MYC SPEARs* carrying the RM9A binding motif forming specific complexes with full-length histone H2A.Z. Right panel: similar REMSA using non-acetylated and acetylated peptides from the N terminal sequence of H2A.Z: (NH_2_-AGGKAGK(Ac)DSGKAKTKAVSRS-COOH). **d**, REMSA gel for *c-MYC SPEARs* carrying the RM9A binding motif and forming specific complexes with TIP60 protein (lane 4) The mutated RNA and DNA motifs do not form complexes (lanes 5 and 6). Both RNA and DNA probes were used at the same molar concentration (0.5 nM), which corresponds to double the counts/minute for the double-stranded DNA duplexes relative to the single-stranded RNA oligonucleotides.

RNA electrophoresis mobility shift assays (REMSAs) using the RM9A motif demonstrated that it is able to form RNPs with H2A.Z and with TIP60 *in vitro* (Fig. 2c, lane 3; and Fig.2d, lane 4). A shift in migration was also observed by incubation of RM9A with synthetic peptides corresponding to the N-terminal sequences of H2A.Z, both unmodified and acetylated at lysine 7 (K7) (Fig. 2c, right panel). Acetylation of the H2A.Z tail peptide did not abrogate the binding (Fig. 2c, lanes12, 13 – acetylated peptide, and lanes 8, 9 – not acetylated peptide), indicating that simple charge-charge interactions are not responsible for *SPEARs*-H2A.Z or *SPEARs*-acH2A.Z complexes. Interestingly, the presence of a predicted stem-loop-like structure (RNAfold^43,44^; Fig. 2e) seems to be required for RM9A binding to H2A.Z and TIP60 (Fig. 2c,d), but unlike the previously identified *DiRs* (DNMT1-interacting RNAs)^45^, the secondary structures are not the only requirement for RNP formation. Indeed, mutation of RM9A so as to prevent the predicted stem-loop structure reduced but did not totally abrogate the formation of the RNA-TIP60 complexes (Fig. 2d, lanes 4-5). Importantly, unrelated oligonucleotides (UR2 and UR3) lacking the common binding motif did not form comparable RNP complexes with TIP60/H2A.Z (Extended Data Fig. 3), indicating the involvement of both primary and secondary structure elements in the recognition of TIP60/H2A.Z by RM9A.

RNAs might be important for binding of remodeling complexes to chromatin, as found for the polycomb repressive complex 2 (PRC2) and Tip60–p400^13,46–48^. To test if *SPEARs* facilitate binding of TIP60 to the chromatin, we compared TIP60 binding to the RNA (single-stranded RM9A) and DNA (double-stranded DM9A) having the same primary sequence. The REMSA/EMSA gels indicate very strong TIP60 binding to the ssRM9A and weaker binding to the dsDM9A (Fig. 2d, lane 4 *vs*. lane 6).

In conclusion, a specific interaction is seen between the RM9A region of the *c-MYC SPEAR* and the histone acetyltransferase TIP60 and histone H2A.Z, in particular its modified form acH2A.Z.

### *SPEARs* are involved in H2A.Z acetylation and exchange

To directly test if *SPEARs* are required for the deposition of acH2A.Z at the TSS of their corresponding gene, transcription was pharmacologically modified and followed by chromatin immunoprecipitation/sequencing (ChIP-PCR and ChIP-Seq) using H2A.Z and acH2A.Z antibodies (outlined on Extended Data Fig. 4a). Two transcription inhibitors, Actinomycin D (ActD; a RNA Polymerase I, II, and III inhibitor) and 5,6-Dichlorobenzimidazole 1-β-D-ribofuranoside (DRB; a RNA Polymerase II Inhibitor), were used at concentrations sufficient to interfere with the transcripts produced by both RNA Polymerases II and III. The rationale behind the use of ActD and DRB is: (i) to globally assess the effect of changes in *SPEARs* expression levels over a short time frame; (ii) to compare effects resulting from the two different pathways triggered by ActD and DRB; and (iii) to take advantage of the reversibility of DRB treatment. Cells were synchronized with a double thymidine block and released into S phase with ActD/DRB added (Extended Data Fig. 4a), treated with the drugs for two hours, an interval during which the overall levels of H2A.Z and TIP60 proteins were not affected (Extended Data Fig. 4b), while the global RNA levels of both the coding genes and the *SPEARs* were significantly altered (Fig. 3a). Relative changes were then investigated in the distributions of unmodified H2A.Z and acH2A.Z in the vicinity of the TSS of the *SPEARs*-linked genes, in response to DRB- and ActD-induced changes in *SPEARs* expression levels. The enrichments of ChIP-Seq signals for H2A.Z and acH2A.Z in the region adjacent to the TSSs of the 2363 *SPEARs*-linked genes following drug treatment were compared with their occupancy in mock-treated (DMSO) cells. We observed substantial reductions in acH2A.Z enrichment at the TSS of samples treated with either DRB or ActD (Fig. 3b, right panels), in contrast to only modest changes in the occupancy levels of unmodified H2A.Z (Fig. 3b, left panels). In cells treated with DRB, 26% of identified *SPEARs* loci showed a drop in acH2A.Z occupancy, while only 6% of such loci showed a similar trend in the H2A.Z signal (Fig. 3b, upper panels). ActD-treated cells showed a more pronounced effect, with 42% of *SPEARs* loci showing reduced acH2A.Z signal as compared with 4% of such loci showing decreased occupancy of H2A.Z (Fig. 3b, bottom panels). Thus, Fig. 3b demonstrates that loci with suppressed expression of *SPEARs* showed diminished enrichment in acH2A.Z, implying that *SPEARs* are involved in the precise placement of this epigenetic mark. The observation of a 10-fold greater number of acH2A.Z-affected TSS in drug-treated samples as compared to H2A.Z, indicates the involvement of *SPEARs* in the maintenance of normal levels of H2A.Z acetylation at TSS, i.e. to the generation of the active epigenetic mark acH2A.Z. The upper panel of Fig. 3c depicts snapshots of the *c-MYC* locus for which the *SPEARs* are repressed by ActD and DRB treatment (results of qRT PCR are shown in Extended Data Fig. 4c, left panel), resulting in a decreased intensity of acH2A.Z peaks. In detail, for the *c-MYC* the ratios of control (DMSO) to drug treated acH2A.Z peak intensities were [DMSO/DRB/ActD]^*c-MYC*^ = 1/0.45/0.39. These ChIP-Seq data were confirmed by quantitative ChIP-qPCR for several amplicons covering the *c-MYC* locus (Extended Data Fig. 4d). By contrast, loci escaping the down-regulating effects of the drugs on their *SPEARs* (the “DRB paradox”^49^) exhibit an acH2A.Z intensity ratio in simple correlation with the change in *SPEARs* levels. For example, the *MYB* locus gives rise to *SPEARs* positively affected by ActD and DRB (Extended Data Fig. 4c, right panel), and exhibits an increase in the intensity of the acH2A.Z peaks, with no significant changes in the intensity of the unmodified H2A.Z peaks (Fig. 3c, bottom panel). For *MYB*, the ratios of control (DMSO) to drug treated acH2A.Z peak intensities are [DMSO/DRB/ActD]^*MYB*^ =1/1.52/1.60, which correlates with the increases in its *SPEARs* levels (Extended Data Fig. 4c, right panel). Thus, loci at which expression of *SPEARs* are positively affected by ActD/DRB illustrate how an increase in *SPEARs* levels leads to a corresponding rise in the intensity of the acH2A.Z peaks.

**Fig. 3.**
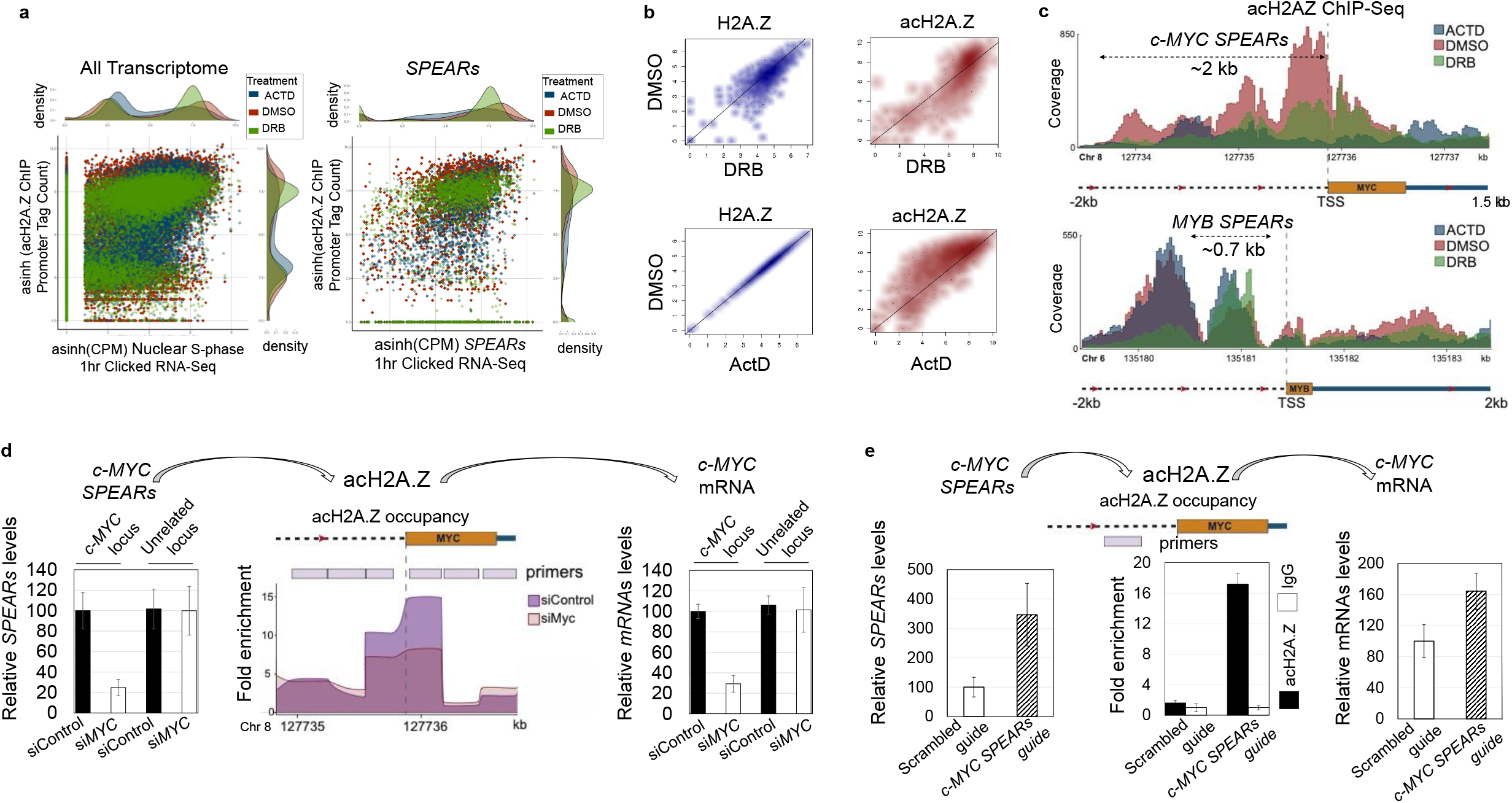
Global and targeted up- and down-regulation of *SPEARs* expression leads to corresponding changes in occupancy of acH2A.Z at the TSSs of linked genes. **a**, Left Panel – Scatterplot of the global inverse hyperbolic sine (asinh) transformed counts-per-million (CPM) normalized Nuclear nascent RNA-Seq expression vs the asinh transformed acH2A.Z ChIP-Seq promoter signal count for DMSO (red), DRB (green), and ActD (blue) treated samples. Right Panel - Scatterplot of the confirmed *SPEARs* asinh transformed counts-per-million (CPM) normalized nuclear nascent RNA-Seq expression vs the asinh transformed acH2A.Z ChIP-Seq promoter signal count for DMSO (red), DRB (green) and ActD (blue) treated samples. Consistent with previous reports, a subset of genes show the “DRB paradox” in which their expression increases post-treatment. **b**, Density plots showing total promoter signal for DRB and ActD treatment vs DMSO for both H2A.Z and acH2A.Z ChIP-Seq samples. Top left panel: DRB treatment showed decreased H2A.Z promoter signal in 6% of genes. Bottom left panel, ActD treatment showed decreased H2A.Z promoter signal in 4% of genes. Top right panel: DRB treatment showed decreased acH2A.Z promoter signal in 26% of genes. Bottom right panel: ActD treatment showed decreased acH2A.Z promoter signal in 42% of genes. **c**, Snapshots of the individual loci, *c-MYC* and *MYB*, for which the *SPEARs* are repressed or induced, respectively, by ActD and DRB treatment (results of qRT PCR are shown in Extended Data Fig. 4c), resulting in corresponding decrease or increase in the intensity of acH2A.Z peaks. **d**, siRNA-induced downregulation of *c-MYC SPEARs* (~75%; left panel) leads: i) to diminished acH2A.Z peaks (middle panel); and ii) to downregulation of *c-MYC mRNA* (~70%; right panel). qRT–PCR, bars indicate mean ±s.d. **e**, CRISPR/dCas9-VP64 gene activation system-based upregulation of the *cMYC SPEARs* (~3-fold; left panel) leads: i) to increased acH2A.Z peaks (middle panel; amplicon located 652 nt to 345 nt from *c-MYC* TSS); and ii) to upregulation of *c-MYC* mRNA (~70%; right panel). The position of tested Guide RNAs is shown in Extended Data Fig. 4f. qRT–PCR, bars indicate mean ±s.d.

Nascent chromatin immunoprecipitation (nasChIP-PCR^50–52^) is particularly useful for tracking histone deposition and was therefore used with the antibodies to H2A.Z, acH2A.Z and TIP60. Cells were synchronized with a double thymidine block, released into S phase and treated with DRB for 2 hours. The medium was supplemented with the EdU DNA analog to enable collection of nascent chromatin DNA (Extended Data Fig. 4a). It was observed that when transcription was inhibited with DRB, the *c-MYC* locus showed diminished enrichment in H2A.Z, acH2A.Z and TIP60 (Extended Data Fig. 4e, three upper panels). This demonstrates a link between suppressed *c-MYC SPEARs* expression and reduced levels of the replacement histone H2A.Z on the nascent chromatin and an even greater reduction of the acetylated acH2A.Z at the TSS of the *c-MYC* locus. Similar results were obtained for the *PU.1* gene locus (Extended Data Fig. 4e, three bottom panels).

Taken together, the negative effects of ActD/DRB – a global “loss-of-function”, exemplified by *c-MYC*, – and their positive effects, a global “gain-of-function”, exemplified by MYB –monitored by ChIP-Seq and nasChIP-PCR experiments, define the role of *SPEARs* transcripts in deposition of the replacement histone H2A.Z. The more robust drop in the acetylation level of H2A.Z as compared to the more modest decrease in the unmodified form, as well as loss of TIP60 at the same sites close to the TSSs of the genes, all point to the role of *SPEARs* transcription in mediating the acetylation of H2A.Z.

In complementary gene-specific experiments, we then tested whether RNAi-mediated downregulation and dCas9-VP64-mediated upregulation of specific *SPEARs* followed by ChIP-Seq and ChIP-qPCR analyses leads to a corresponding reduction or increase of H2A.Z and acH2A.Z levels at the TSS of targeted loci and in reduced or increased expression of the corresponding gene. Knockdown of the *c-MYC SPEARs* (~75%; Fig. 3d, left panel) was associated with a significant decrease of acH2A.Z levels at the TSS of the *c-MYC* gene (Fig. 3d, middle panel), followed by the corresponding decrease of *c-MYC* mRNA expression of similar magnitude (~70%; Fig. 3d, right panel). The results of the RNAi/ChIP knockdown experiments demonstrate that reduced expression of the *c-MYC SPEARs* is linked to lower levels of H2A.Z acetylation at the *c-MYC* TSS, which accords with the initial hypothesis of direct causality between the expression of *SPEARs* and the deposition of the activating histone mark, leading to transcriptional activation of the adjacent gene.

Additionally, we performed a “gain-of-function” test using the dCas9-VP64 gene activation system (outlined on Extended Data Fig. 4f). Increasing levels of *c-MYC SPEARs* expression (Fig. 3e, left panel) were associated with a significant increase of acH2A.Z levels at the TSS of the *c-MYC* gene (Fig. 3e, middle panel) accompanied by a ~2-fold increase in *c-MYC* gene expression (Fig. 3e, right panel).

Overall, downregulation of *SPEARs* using independent pharmacological and RNAi-induced methods leads to a substantial reduction in the activating epigenetic mark acH2A.Z. Loss of the acetylated form of H2A.Z from chromatin in the vicinity of active TSS implies a reduction of its activity, i.e. reduced expression of the corresponding gene (Fig. 3d, right panel). On the other hand, upregulation of *SPEARs* leads to a substantial induction of the activating epigenetic mark acH2A.Z and, consequently, to elevated expression of the corresponding gene (Fig. 3e, right panel).

### *SPEARs* regulate the expression of the corresponding mRNA *via* TIP60 recruitment and acH2A.Z deposition

To demonstrate that the *SPEARs*-mediated regulation of their corresponding mRNA is executed through the TIP60/acH2A.Z pathway, the effects of TIP60 inhibition on the *c-MYC* locus were tested. Two TIP60/HAT inhibitors, TH1834^53^ and MG-149^54^, were used in HL-60 cells to reduce the total level of H2A.Z acetylation (Extended Data Fig. 5a). This led to significant downregulation of mRNAs, in contrast to almost unchanged levels of *SPEARs* transcripts (Fig. 4a). To examine the direct regulation of *c-MYC* expression by TIP60 and acH2A.Z interacting with the *SPEARs*, we monitored the regulating effect of nascent *SPEARs* on the level of nascent mRNA expression in the presence or absence of the two TIP60 inhibitors, taking advantage of the reversibility of the DRB transcriptional inhibitor (ref. ^55^. The rationale was to analyze only transcripts re-appearing after release from the transcriptional block whilst the HAT is inhibited. The short time of inhibitor treatment should rule out secondary effects of HAT inhibition.

**Fig. 4.**
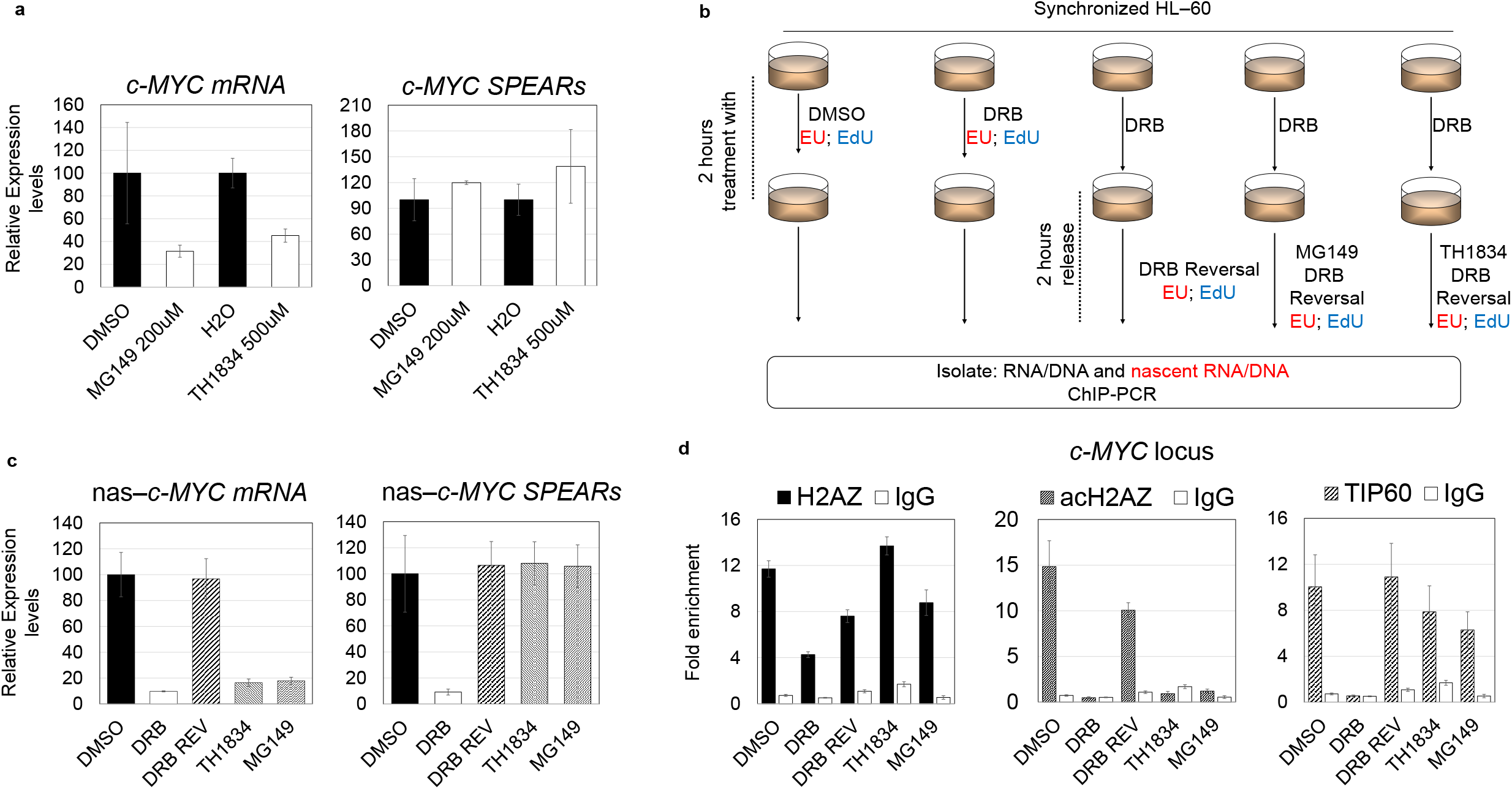
*SPEARs* regulate the expression of their linked mRNA *via* a TIP60/acH2A.Z pathway. **a**, Response of total *c-MYC* mRNA and *SPEARs* to TIP60/HAT inhibitors. *C-MYC* mRNAs are downregulated by MG149 and TH1834, but these inhibitors do not affect *c-MYC SPEARs*. HL-60 cells were released from double thymidine block and treated for 2 hours with MG149 (200 μM) and TH1834 (500 μM); control (mock) treatments were supplemented with DMSO (0.05%) or water, respectively. qRT–PCR and strand-specific qRT-PCR were used to quantitate *SPEARs* and mRNA. Bars indicate mean ±s.d. **b**, Experimental design: HL-60 cells were released into S Phase and treated with DMSO/DRB for 2 hrs (as described for Panel A), then washed and incubated for another 2 hrs with or without TIP60 inhibitors. The medium was supplemented with the EU RNA and EdU DNA analogs to enable collection of nascent RNAs and DNAs. Chromatin was collected for ChIP assays with antibodies to H2A.Z, acH2A.Z, TIP60, and IgG. Nascent DNAs and RNAs were isolated from the immunoprecipitated chromatin and from total RNA, respectively (see Methods for details; Extended Data Fig. 1a) and analyzed by qPCR and strand-specific qRT-PCR, respectively. **c**, Different response of nascent *c-MYC SPEARs* and *c-MYC* mRNA to TIP60/HAT inhibitors: *c-MYC* mRNAs are downregulated but *c-MYC SPEARs* are not affected. Quantitation was performed by qRT-PCR (for mRNA) and strand-specific qRT-PCR (for *SPEARs*): bars indicate mean ±s.d. **d**, Nascent ChIP-qPCRs for the *c-MYC* locus using antibodies to: H2A.Z, acH2A.Z, and TIP60 with an IgG control (amplicon located 652 nt to 345 nt from *c-MYC* TSS). These demonstrate: (i) DRB treatment (which leads to downregulation of *c-MYC SPEARs*) results in the loss of acH2A.Z (middle panel) and TIP60 (right panel) and a partial loss of H2A.Z (left panel); (ii) Restoration of *c-MYC SPEARs* (following DRB reversal) leads to reappearance of H2A.Z, acH2A.Z and TIP60; and (iii) Inhibition of TIP60 activity by MG149 and TH1834 prevents the restoration of acH2A.Z, while only slightly affecting = TIP60 and H2A.Z enrichment. Quantitation by qPCR: bars indicate mean ± s.d. (n=2).

Cells were synchronized with a double thymidine block and released into S Phase with DMSO/DRB added. After 2 hours (during which H2A.Z levels remained unaffected, but *SPEARs* and mRNA levels drop (see Extended Data Fig. 5a,b), cells were washed and then incubated for a further 2 hours with or without the TIP60 inhibitors. This interval is sufficient for the levels of both the *c-MYC* mRNAs and its *SPEARs* to be restored to the levels when the TIP60 inhibitors are absent (results shown in Extended Data Fig. 5b). The medium was supplemented with the EU RNA and EdU DNA analogs to enable collection of nascent RNAs and newly formed chromatin that had escaped the drug-induced inhibition of transcription or acetylation. After 2 hours with the TIP60 inhibitors present (MG-149 at 200 μM and TH1834 at 500 μM), during which time the levels of acH2A.Z dropped (see Extended Data Fig. 5a), cells were crosslinked and subjected to nasChIP-PCR and nascent RNA expression analyses (nas-qRT-PCR) (outlined on Fig. 4b). Collected RNAs were biotinylated by click chemistry, isolated on streptavidin beads and analyzed by nas-qRT-PCR (see Methods for details and Extended Data Fig. 1a, bottom panel). The results in Fig. 4c show that the correlation between expression of the *SPEARs* and the mRNA no longer holds when TIP60 activity is inhibited, *i.e*. the restored levels of the *SPEARs* are incapable of rescuing the expression of *c-MYC* mRNA to the level defined by the reversal of DRB repression. These data suggest that the function of *SPEARs* in regulating *c-MYC* mRNA expression is impeded by inhibition of the HAT activity of TIP60.

To further check that *SPEARs* realize their *c-MYC* regulatory function directly through the TIP60/acH2A.Z pathway, we assessed the differences in chromatin occupancy of H2A.Z, acH2A.Z and TIP60 after inhibition of transcription (“DRB”), reversal of inhibition (“DRB REV”), and reversal of inhibition in the presence of the two TIP60 inhibitors, using nasChIP-qPCR. In particular, it was important to check whether the drop in *c-MYC* expression after treating the cells with DRB and TIP60 inhibitors is correlated with diminished enrichment in TIP60, acH2A.Z, or H2A.Z. Nascent DNA was isolated from the chromatin immuno-precipitated with antibodies to H2A.Z, acH2A.Z, or TIP60, biotinylated by click chemistry, isolated on streptavidin beads (Methods) and finally analyzed by qPCR at amplicons corresponding to maximum enrichment within the *c-MYC* locus (see Fig. 3e). By analyzing the nascent RNAs and chromatin, we examined only the immediate changes in *c-MYC* expression and TIP60, acH2A.Z, and H2A.Z occupancy imposed by the inhibition of transcription and/or the reduced acetylation of histones. The middle and right panels of Fig. 4d demonstrate that the *c-MYC* locus with suppressed expression of its *SPEARs* shows diminished enrichment of both acH2A.Z and TIP60, and the reversal of the transcription inhibition led to the restoration of both enrichments. This indicates that the *SPEARs* are involved in the recruitment of TIP60 and in the proper placement of the acetylation mark. When the two HAT inhibitors are present, the reversal of transcriptional inhibition can only restore the enrichment levels of unmodified H2A.Z and of TIP60 (Fig. 4d, left and right panels) but not the levels of acH2A.Z (Fig. 4d, middle panel). In contrast, no enrichments of H2A.Z, acH2A.Z, or TIP60 were detected within the Gene Desert locus^56^ (Extended Data Fig. 5c). Given that local levels (i.e. in the vicinity of the *c-MYC locus* TSS; Fig. 4d, left panel) as well as overall levels of H2A.Z were unchanged throughout the experiment (Extended Data Fig. 5a), these results establish the leading role of the TIP60/acH2A.Z pathway in *c-MYC SPEARs*-mediated regulation of its corresponding gene, *c-MYC*. Similar results were obtained for the *PU.1* locus (Extended Data Fig. 5d,e) and with another HAT inhibitor – NU 9056 (which at high concentration inhibits acetylation of H2A.Z) (Extended Data Fig. 5f-h). As with HAT inhibitors MG149 and TH1834 (Fig. 4d, middle panel), we observed loss of promoter occupancy by acH2A.Z after treatment with the HAT inhibitor NU 9056 in cells pre-treated with DRB (“DRB/NU 9056”) (Extended Data Fig. 5h).

Collectively our results suggest that the TIP60/acH2A.Z pathway is a general mechanism for *SPEARs*-mediated regulation of the corresponding adjacent coding genes.

### *SPEARs* regulate recycling of the epigenetic mark - acH2A.Z

To demonstrate the role of *SPEARs* in the recycling of the acH2A.Z mark, we performed the experiments outlined in Fig. 5a. Cells were cultured for six days in medium supplemented with “heavy” ^15^N_2_^13^C_2_-Lysine and ^13^C_6_-D-glucose and then synchronized with a double thymidine block. After washing out “heavy” lysine and glucose, cells were released into S phase in the presence of HAT inhibitor or DMSO (mock treatment). The rationale of the experiment was to distinguish histones deposited in the chromatin before entering S phase from those deposited after this point. Thus, histones containing “heavy” lysines and their acetylated forms carrying “heavy” acetyl groups are “old”, whilst “new” histones and their acetylated forms generated during early S phase should contain “light” lysines and “light” acetyl groups. The medium was supplemented with the EdU DNA analogs to enable collection of newly formed (nascent) chromatin. The HAT inhibitor NU 9056 was added at the start of S-phase and the efficiency of inhibition was demonstrated by HAT assays on the collected cells (Extended Data Fig. 6a) showing efficient suppression of H2A.Z acetylation. This experiment enabled investigation of whether “old” histones with “old” modifications are preserved and transmitted through the generational border. At 4 hours after release from synchrony, cells were collected and subjected to gene expression and Western Blot analyses, HAT assay, ChIP-PCR, nasChIP-PCR, and mass spectrometry. As with TH1834 and MG148 inhibition (Fig. 5a,b), treatment with NU 9056 resulted in a drop in mRNA but not in *SPEARs* expression (Fig. 5b) and levels of H2A.Z remained unaffected but acH2A.Z levels dropped (Fig. 5c).

**Fig. 5.**
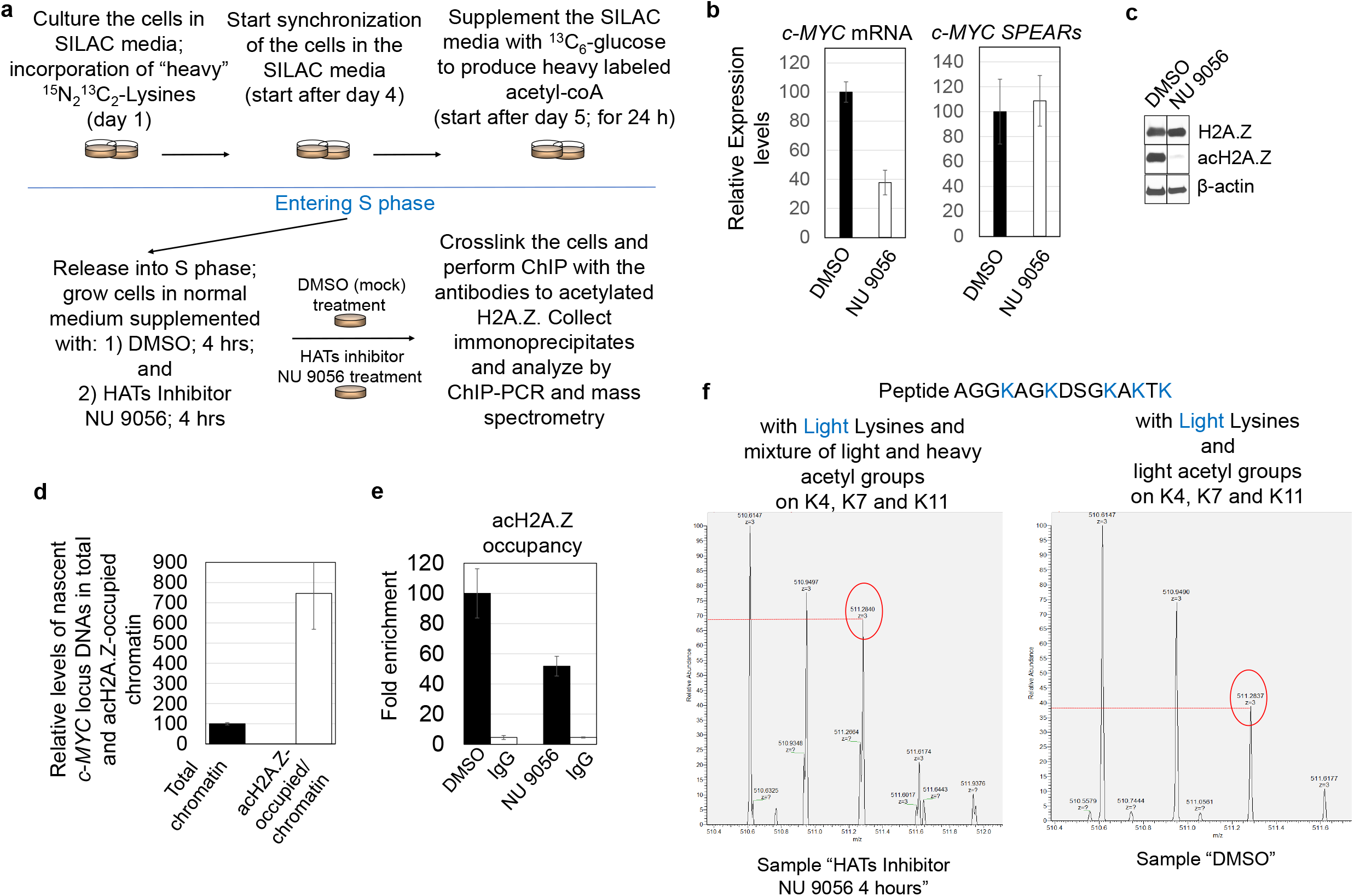
*SPEARs* regulate the recycling of the epigenetic acetylation mark on histone H2A.Z. **a**, Outline of experimental design. **b**, Response of total *c-MYC* mRNA and *SPEARs* to the treatment with TIP60/HAT inhibitor NU 9056. **c**, Western blot analyses of proteins isolated 4 hours after release from synchrony and treatment with TIP60/HAT inhibitor NU 9056, showing that the global content of H2A.Z is not affected by HAT-inhibitor treatment, in contrast to acH2A.Z, which is highly depleted, i.e. the modification is largely absent. **d**, Nascent ChIP-qPCRs monitored at the peak of the *c-MYC* locus using antibodies to acH2A.Z showing a 7-fold enrichment over total chromatin. **e**, ChIP-qPCRs for the *c-MYC* locus using antibodies to acH2A.Z following treatment with HAT inhibitor NU 9056; (amplicon located 652 nt to 345 nt upstream of the *c-MYC* TSS). **f**, Examples of the detected N-terminal peptides from sample “HAT inhibitor (NU 9056)-treated sample (4 hours)”. Left Panel: Peptides with all light Lysines and light or heavy acetyl groups (the latter carry the ^13^C_6_ carbon atoms and add height toward the third isotopic peak in the distribution). When compared to the same N-terminal peptide with no heavy acetyl groups, shown in the RH panel and representing natural abundance of the third isotope in the distribution, the heavy acetyl groups from HAT inhibitor treated sample shows elevated height of the third peak from 37% of natural distribution (compare to the first ^12^C peak) to about 67%. Observed change in the third isotope abundance between the two samples shows that in the HAT inhibited sample about 55% of the N-terminal peptide contain heavy acetyl groups (37%/67% x 100% = 55%). Right Panel: Mock (DMSO)-treated sample shows the N-terminal peptide (AGGKAGKDSGKAKTK) with three heavy Lysines and the absence of incorporated heavy acetyl groups. The HAT inhibitor (NU9056)-treated sample (4 hrs) shows that the ratio of the N-terminal peptide with three light Lysines (at m/z ~510) *vs*. the same peptide with all three heavy Lysines (at m/z ~524) was measured as ~1.8 (see Extended Data Fig. 6c)

To verify that the *c-MYC* locus indeed undergoes replication and chromatin re-assembly in the early part of S phase, nasChIP-PCR was performed (with antibodies to acH2A.Z) and a significant enrichment of acH2A.Z-occupied nascent *c-MYC* locus chromatin was observed (Fig. 5d).

The tests of experimental conditions being satisfactory (Extended Data Fig. 6a and Supplementary Data #5), ChIP-PCR analyses of acH2A.Z occupancy at the target promoters was performed. Strikingly, analysis of promoter occupancy by acH2A.Z in HL-60 cells after even longer treatment in NU 9056 showed levels of acH2A.Z to be diminished by only one half in comparison to the mock treatment (Fig. 5e, right panel). Two points need emphasis: (i) when acetylation of all newly synthesized histones is strongly inhibited, persistent promoter occupancy by the acetylated forms of H2A.Z can occur *only* through the recycling of the previously acetylated (“old”) histones; and (ii) this recycling process is interrupted by the inhibition of transcription, particularly with the abrogation of *SPEARs* expression (see Fig. 4 and 5).

To unequivocally demonstrate that the “recycling” of acetylated forms of histones indeed represents “old” histones, we performed mass spectrometry analyses of the immunoprecipitates from cells subjected to HAT inhibitor for 4 hours” (Fig. 5a), in which all “new” acetylation was suppressed. The samples were digested with chymotrypsin and analyzed on an Orbitrap Elite mass spectrometer. The Proteome Discoverer 2.4 search engine analyzed the raw data. Searches were performed against the H2A.Z protein sequence and common contaminants and all results filtered to 1 % FDR (false discover rate). The analysis focused on the N-terminal H2A.Z peptide Ac-AGGKAGKDSGKAKTK (Extended Data Fig. 6b) with variable acetylation of lysines K4, K7, and K11 and showed a high level of “heavy” lysines and the presence of heavy acetyl groups in this N-terminal peptide. Analysis of isotopic distributions showed that among all N-terminal peptides carrying two acetyl groups, the content of heavy lysines was ~70% and of heavy acetyl groups was ~20%. In peptides carrying three acetyl groups the content of heavy lysines was ~55% and of heavy acetyl groups ~30%. Extended Data Fig. 6c shows examples of the peptides detected. Fig. 5f demonstrates an increase of heavy acetyl groups in the “HATs inhibitor 4 hours” sample (Left panel; ~70%) as compared to their natural distribution (Right panel; ~38%). These analyses show that some old H2A.Z histones carrying heavy acetyl groups are preserved in the early stages of S phase and that recycling of old acH2A.Z is affected by brief suppression of transcription in the first hours of S phase.

## Discussion

The roles of ncRNAs in regulating deposition and recycling of acH2A.Z at gene promoters has been investigated. Transcription of *SPEARs* that control the acetylation of nearby H2A.Z could be regarded as a mechanism whereby information encoded in the genome becomes expressed as ‘epigenetic’ information. Locally induced *SPEARs* bind to the replacement histone H2A.Z and to a nuclear factor, the histone acetyl transferase TIP60, leading to deposition/acetylation of H2A.Z, a process that represents recycling of the mark across generations. *SPEARs* transcripts guide deposition/acetylation of this replacement histone through RNP formation with TIP60/H2AZ. The function of these transcripts depends on a combination of their expression levels, primary and secondary structures, and physical co-compartmentalization of RNAs, their parental loci, and their protein partners. *SPEARs* databases can provide a valuable resource for studying the dynamics of RNP assembly and their role in the formation of active/inactive chromatin. This mechanistic model is depicted in Figure 6.

**Fig. 6.**
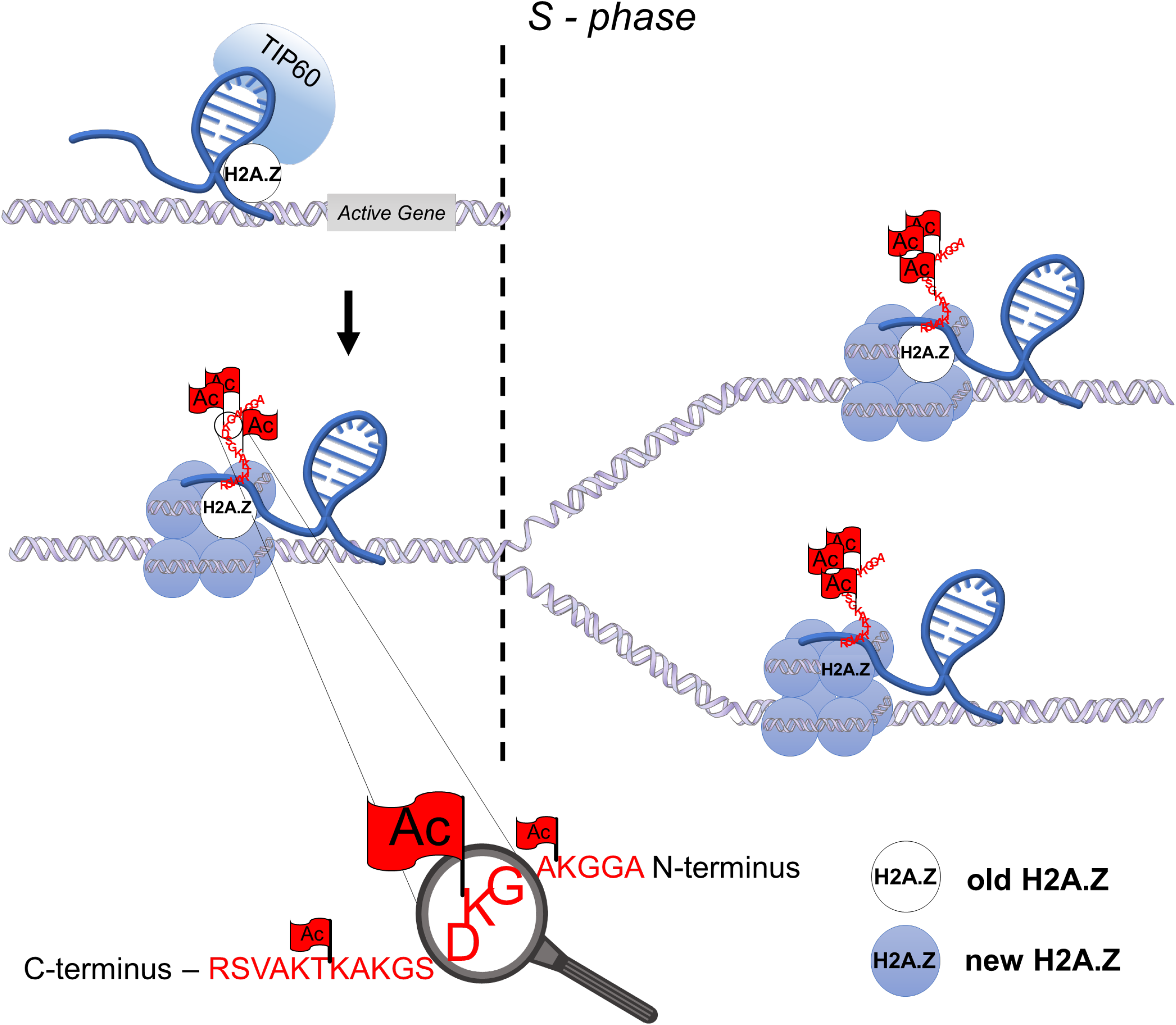
Model: During early S phase, *SPEARs* interact with both histone H2A.Z and the acetyltransferase TIP60. H2A.Z and TIP60 achieve physical proximity, leading to a high local effective protein concentration^60^ that favors H2A.Z acetylation and exchange with canonical H2A within the nucleosome. In this active chromatin conformation, gene expression is initiated. The incorporated “old” acetylated H2A.Z marks are preserved and transmitted through the generational border.

The demonstrated link between *SPEARs* and the activated form of the replacement histone H2A.Z raises the possibility that other long ncRNAs (lncRNAs) play a role in the recruitment of chromatin modifiers that deposit activating or repressive epigenetic marks. Although more detail is still needed, such lncRNAs may act not only as signaling molecules but also as the scaffolds around which chromatin writer/eraser complexes are assembled. This notion is supported by an observation in *Drosophila*, in which the deposition of the replacement histone H3.3 occurs within transcriptionally active loci and is not replication-coupled^57^.

In summary, the discovery of *SPEARs* reveals a novel mechanism for the biosynthesis and maintenance of a major epigenetic mark. Taken together with the study of identified *DiRs*^45^, the present investigations demonstrate that epigenetics and genetics are intrinsically linked, and such lncRNAs modulate distinct epigenetic marks by using site-specific targeting molecules. Indeed, epigenetic modulating agents are already employed in the clinic^58,59^, but assessment of their function is difficult, their efficacy being compromised by lack of specificity. This research provides a new tool to overcome these drawbacks by using the specificity of RNA molecules to drive deposition and selective modification of histones. This study indicates the need for further investigations uncovering how epigenetic information is encoded in the genome and regulated under normal conditions and in disease states.

## METHODS

Detailed methods are provided in the online version of this paper and include the following:

- KEY RESOURCES
- CONTACT FOR REAGENT AND RESOURCE SHARING
- EXPERIMENTAL MODEL
  Mammalian Cell Culture
- METHOD DETAILS
  RNA isolation;
  qRT-PCR;
  Nascent RNA/DNA capture;
  Primer extension and 5’/3’ RACE;
  Double Thymidine block (early S-phase block);
  DRB and Actinomycin D treatments;
  Down-regulation of *c-MYC SPEARs*;
  Ribonucleoprotein (RNP) fractionation;
  Western Blotting Analysis;
  Tandem Mass Spectrometry (LC-MS/MS);
  Nuclear Chromatin (ChIP) and RNA immunoprecipitation (nRIP);
  RNA electrophoretic gel mobility shift assays (REMSAs).
- QUANTIFICATION AND STATISTICAL ANALYSIS
  ChIP- Sequencing Analyses;
  Comparison of the enrichment of H2A.Z and acH2A.Z ChIP-Seq signal surrounding gene TSS loci (+/- 2kb);
  RIP-Sequencing Analyses;
  RNA-Sequencing Analyses;
  Motif Discovery;
  LC-MS/MS Analyses.
- DATA AND SOFTWARE AVAILABILITY
  Sequencing Data are available on the gene omnibus database under the accession ID number: GSE165526; enter token: slehymyczncvfqd. List of all data sets is in Supplementary Data #6.
  Mass Spectrometry data will be deposited to the ProteomeXChange Consortium via PRIDE upon acceptance of the manuscript. Private partial submission with MASSIVE: https://massive.ucsd.edu/ProteoSAFe/dataset.jsp?task=38a5eae58dd247f5b3c42f2be181e692; Password: Alex_Acetyl

## Acknowledgements

This work was supported by the award of NCI R35 CA197697, R01DK103858, W81XWH-15-1-0161, and P01HL131477-01A1 to DGT; and R50 CA211304 to AKE; ADR was supported by NCI R00 CA188595, W81XWH-20-1-0518, the Italian Association for Cancer Research (AIRC) and the Giovanni Armenise-Harvard Foundation; AW was supported by the Belgian American Educational Foundation (BAEF) and by Wallonie-Bruxelles International (WBI); K01CA222707 was awarded to BQT; TB was supported by the National Research Foundation and the Singapore Ministry of Education under its Centres of Excellence initiative. This work was also supported by the Singapore Ministry of Health’s National Medical Research Council under its Singapore Translational Research (STaR) Investigator Award; by the National Research Foundation Singapore and the Singapore Ministry of Education under its Research Centres of Excellence initiative; and by the Singapore Ministry of Education Academic Research Fund Tier 3, grant number MOE2014-T3-1-006. We thank Bruno Amati for thorough discussion and insightful comments on the manuscript and for providing the TIP60 antibodies; likewise Pier Paolo Pandolfi for critical reading of the manuscript and all the members of the Tenen Laboratory for the helpful suggestions.

## Author Contributions

AKE, ADR, CCR and DGT conceived and designed the study and wrote the manuscript; AKE, SU, HZ, ADR, LY, DT, SU, EM, AW, BQT, RC, MRS, HZ, YZ performed experiments; MAB performed bioinformatics and statistical analyses, and analyzed RNA-Seq and RIP-Seq data; VEA, RTM, MAB and TB analyzed ChIP-Seq data, EM and VP performed the RNA motif discovery, BB and DK analyzed Mass Spectrometry datasets.

