## Supplementary material for "Formation and recycling of an active epigenetic mark mediated by cell cycle-specific RNAs": Suppl. Figures

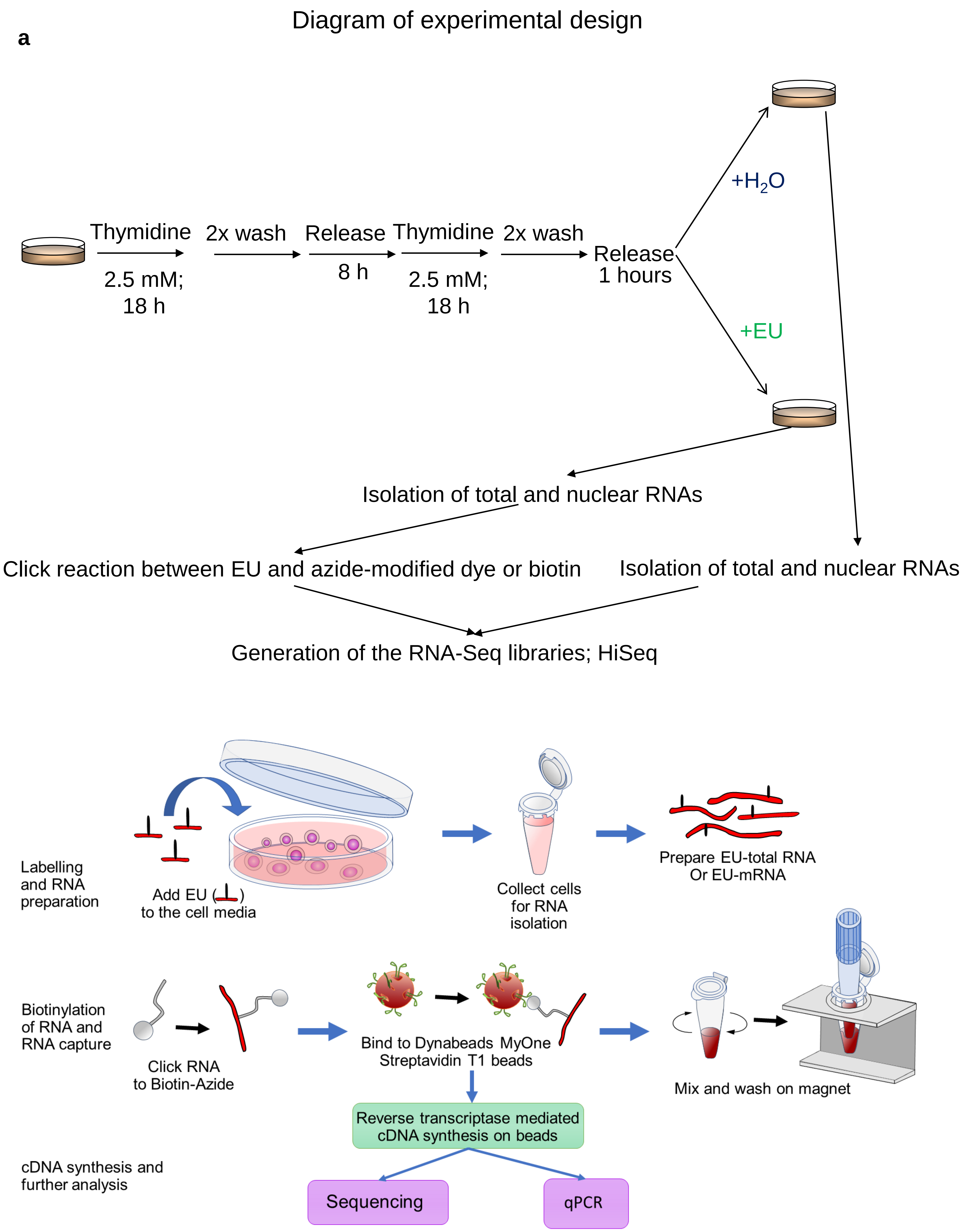


Ebralidze *et.* Extended Data Fig.1_part 1

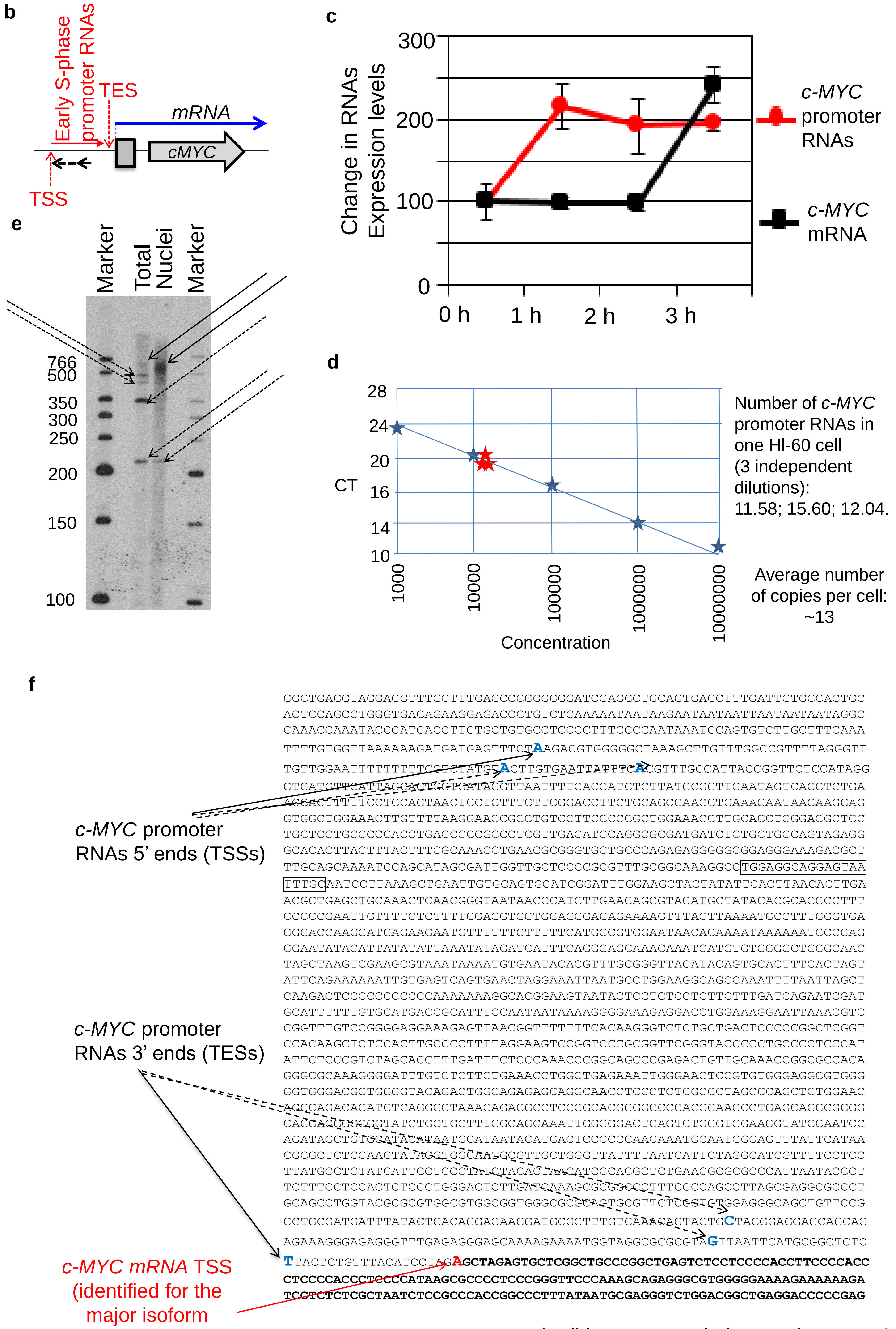


Ebralidze *et.* Extended Data Fig.1_part 2

**Extended Data Fig. 1. Identification of the S phase Early Promoter RNAs.**

**a**, Schematic diagram showing synchronization of HL-60 cells by double thymidine block. Upon release from double thymidine block, cells were spiked with analog EU for downstream Click-iT conversion. After 1 hour, total and nuclei RNA were collected. RNAs were processed according to the manufacturer's recommendation and RNA-seq libraries were generated, sequenced and analyzed. **b**, Diagram of *c-MYC* locus transcripts. Vertical arrows indicate transcription start sites (TSS) and transcription end sites (TES), dashed arrow indicate position of primer in Primer extension experiment. **c**, Levels of coding (mRNA) and promoter RNA immediately after release from double thymidine block. Induction of the *c-MYC* promoter RNAs preceded and exceeded expression of mRNA (qRT-PCR; Bars indicate mean ± s.d.). **d**, *c-MYC* promoter RNAs copy number (Experimental details are available in STAR Methods). **e**, Primer extension experiments. Shown is the radio autograph of the primer extension reactions for the *c-MYC* promoter RNAs performed on total cellular and total nuclei. Black arrows indicate the longest extension product. Dashed arrows indicate "strong-stops" of the extension reactions (position of primer is shown in Figure S1F, boxed sequence; Experimental details are available in STAR Methods). **f**, 5'3' RACE. The "longest" isoform of the early promoter RNAs: TSS at -2160 nt to *c-MYC* mRNA TSS; and TES at -38 nt to *c-MYC* mRNA TSS (nucleotide positions marked by the dark arrows). TSSs and TESs for the "shorter" isoforms are indicated by the dashed arrows (Experimental details are available in STAR Methods).

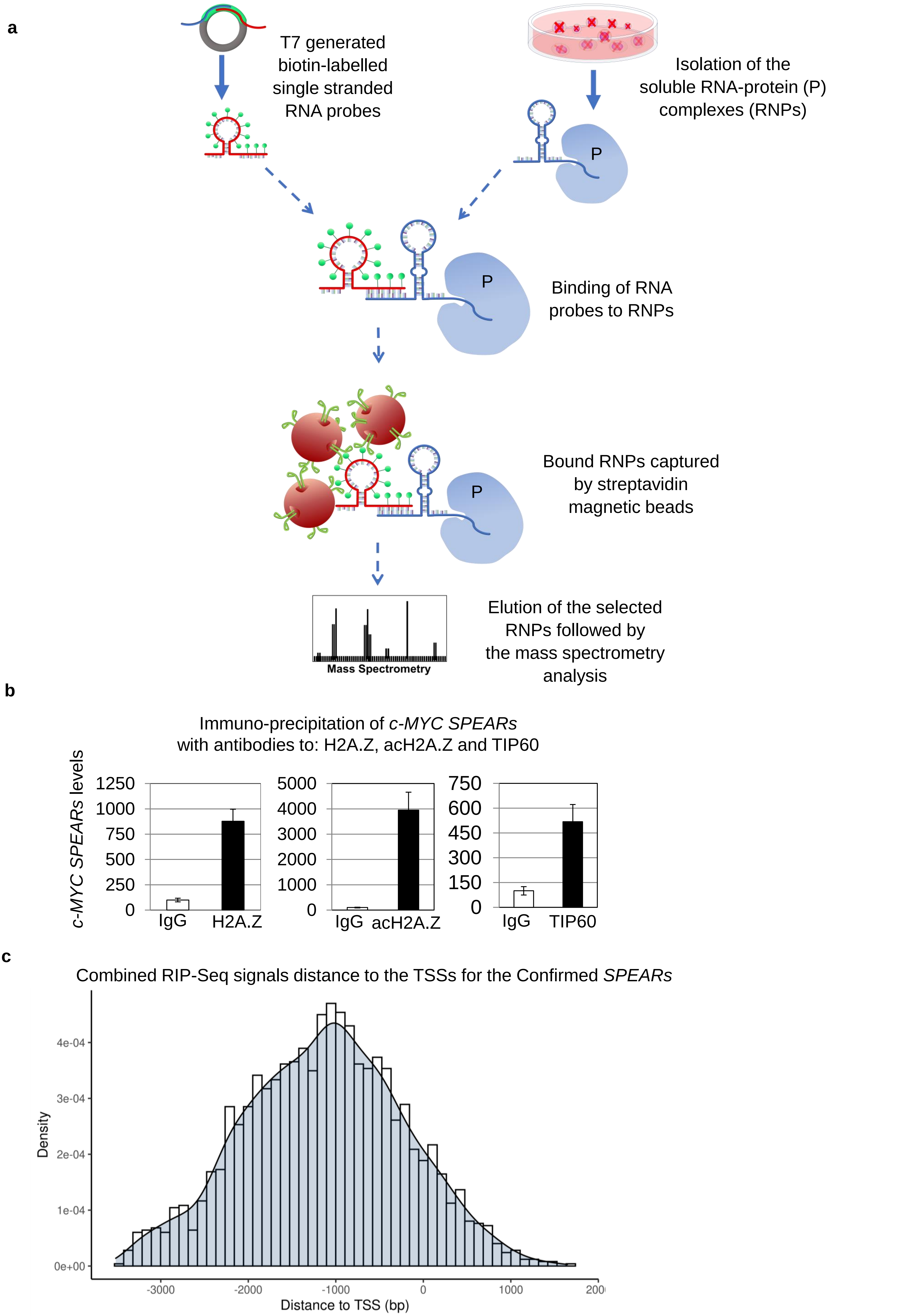
a
T7 generated
biotin-labelled
single stranded
RNA probes
Isolation of the
soluble RNA-protein (P)
complexes (RNPs)
P
P
Binding of RNA
probes to RNPs
P
Bound RNPs captured
by streptavidin
magnetic beads
Mass Spectrometry
Elution of the selected
RNPs followed by
the mass spectrometry
analysis
b
Immuno-precipitation of *c-MYC SPEARs*
with antibodies to: H2A.Z, acH2A.Z and TIP60
*c-MYC SPEARs* levels
1250
1000
750
500
250
0
IgG
H2A.Z
5000
4000
3000
2000
1000
0
IgG
acH2A.Z
750
600
450
300
150
0
IgG
TIP60
c
Combined RIP-Seq signals distance to the TSSs for the Confirmed *SPEARs*
Density
4e-04
3e-04
2e-04
1e-04
0e+00
-3000
-2000
-1000
0
1000
200
Distance to TSS (bp)

**Extended Data Fig. 2. *SPEARs*-H2A.Z-acH2A.Z-TIP 60 interactions and immunoprecipitation.**

**a**, Diagram representing generation of biotinylated *SPEARs* probes and protocol used to pull-down *SPEARs*-containing RNA-protein complexes (*SPEARs*-RNPs). Collected soluble *SPEARs*-RNPs were separated on the 5% PAGE and submitted for Mass spectrometry analyses (Experimental details are available in STAR Methods). **b**, *c-MYC SPEARs* are immunoprecipitated with anti-H2A.Z, -acH2A.Z and -TIP60 antibodies (qRT–PCR, bars indicate mean +/-s.d.). **c**, Assessment of the confirmed *SPEARs* binding location with respect to the TSS shows a normal distribution of distances, with *SPEARs* binding preferentially -1kb upstream of the TSS.

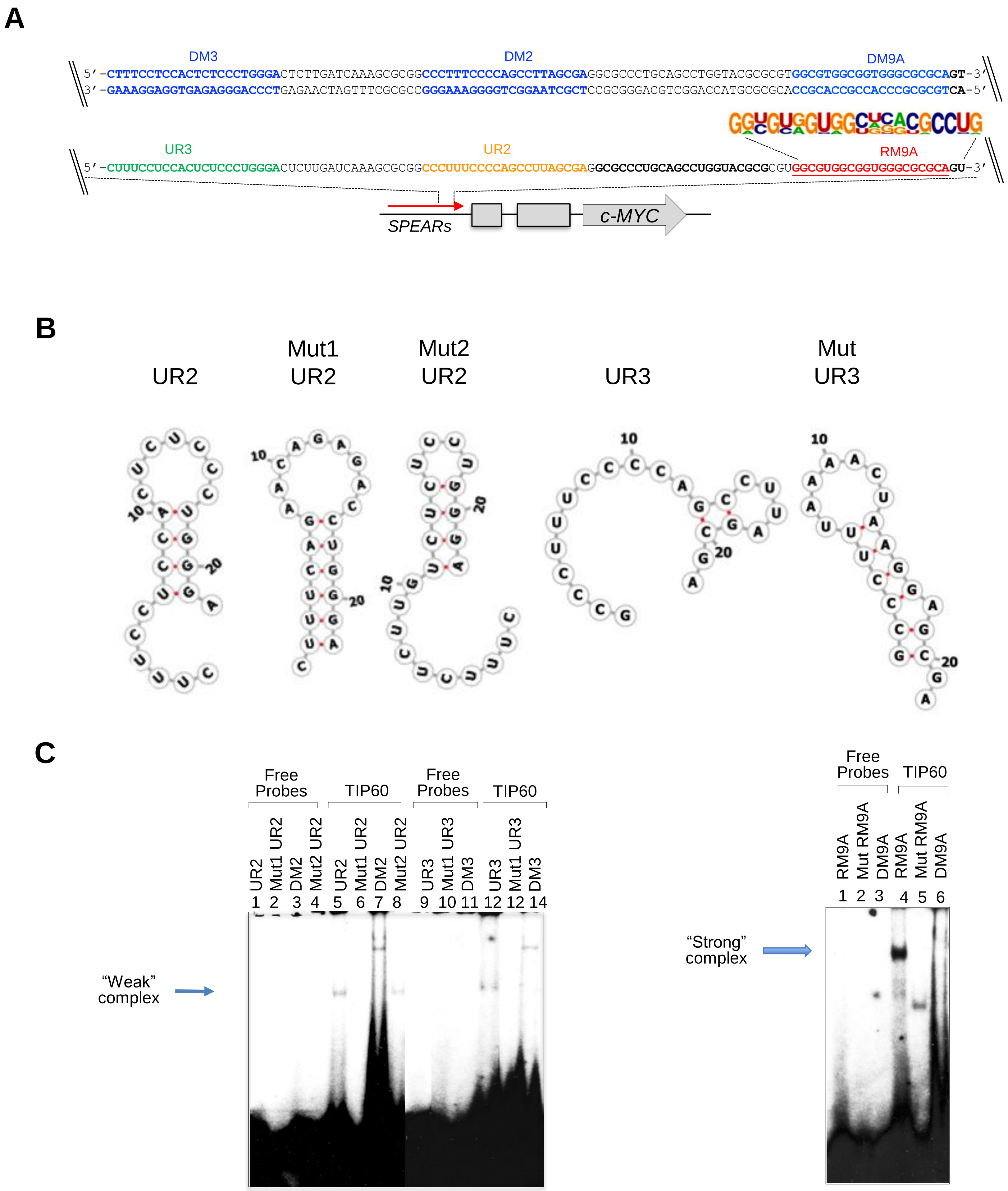


**Extended Data Fig. 3. *SPEARs*-H2A.Z-acH2A.Z-TIP 60 interactions *in vitro*. Identification of common binding motifs in *SPEARs*.**

**a**, The sequence of the common binding motif "9" (RNA oligonucleotide RM9A), the unrelated RNA oligonucleotides 2 (UR2) and 3 (UR3) within the *c-MYC SPEARs* and the corresponding DNA duplexes (DM) are shown. **b**, The RNA secondary structures predicted by RNAfold[43,44] for ribonucleotides used in REMSA experiments. **c**, Left Panel: Unrelated oligonucleotides (UR2 and UR3) lacking the common binding motif did not form comparable RNPs complexes with TIP60. Right Panel: Reproduction of Figure 3D from the main text.

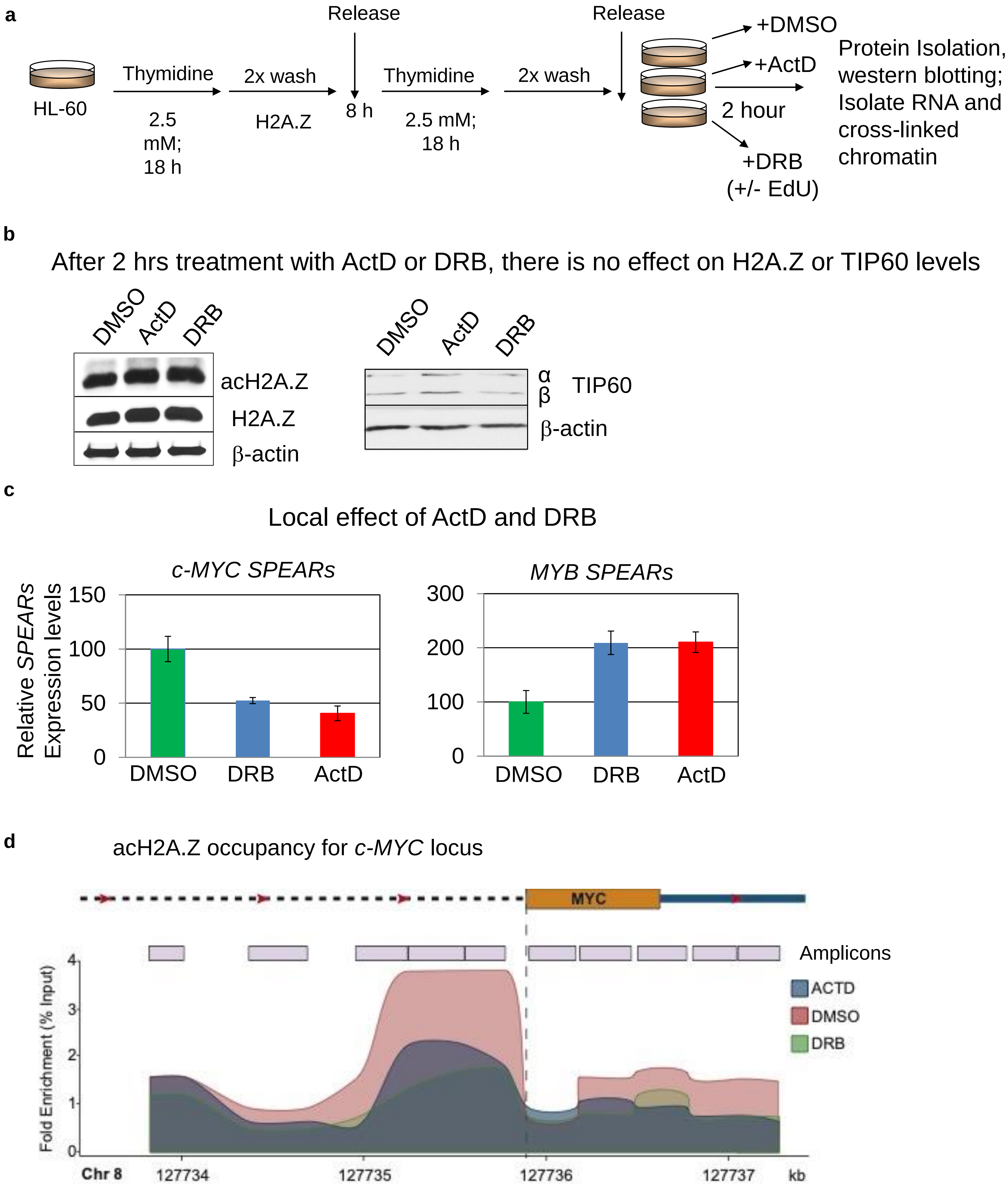
a
HL-60
Thymidine
2.5 mM; 18 h
2x wash
H2A.Z
Release
8 h
Thymidine
2.5 mM; 18 h
2x wash
Release
+DMSO
+ActD
2 hour
+DRB (+/- EdU)
Protein Isolation, western blotting; Isolate RNA and cross-linked chromatin
b
After 2 hrs treatment with ActD or DRB, there is no effect on H2A.Z or TIP60 levels
DMSO
ActD
DRB
acH2A.Z
H2A.Z
β-actin
α
β
TIP60
β-actin
c
Local effect of ActD and DRB
c-MYC SPEARs
MYB SPEARs
Relative SPEARs Expression levels
150
100
50
0
300
200
100
0
DMSO
DRB
ActD
d
acH2A.Z occupancy for c-MYC locus
MYC
Amplicons
ACTD
DMSO
DRB
Fold Enrichment (% Input)
4
3
2
1
0
Chr 8
127734
127735
127736
127737
kb

e

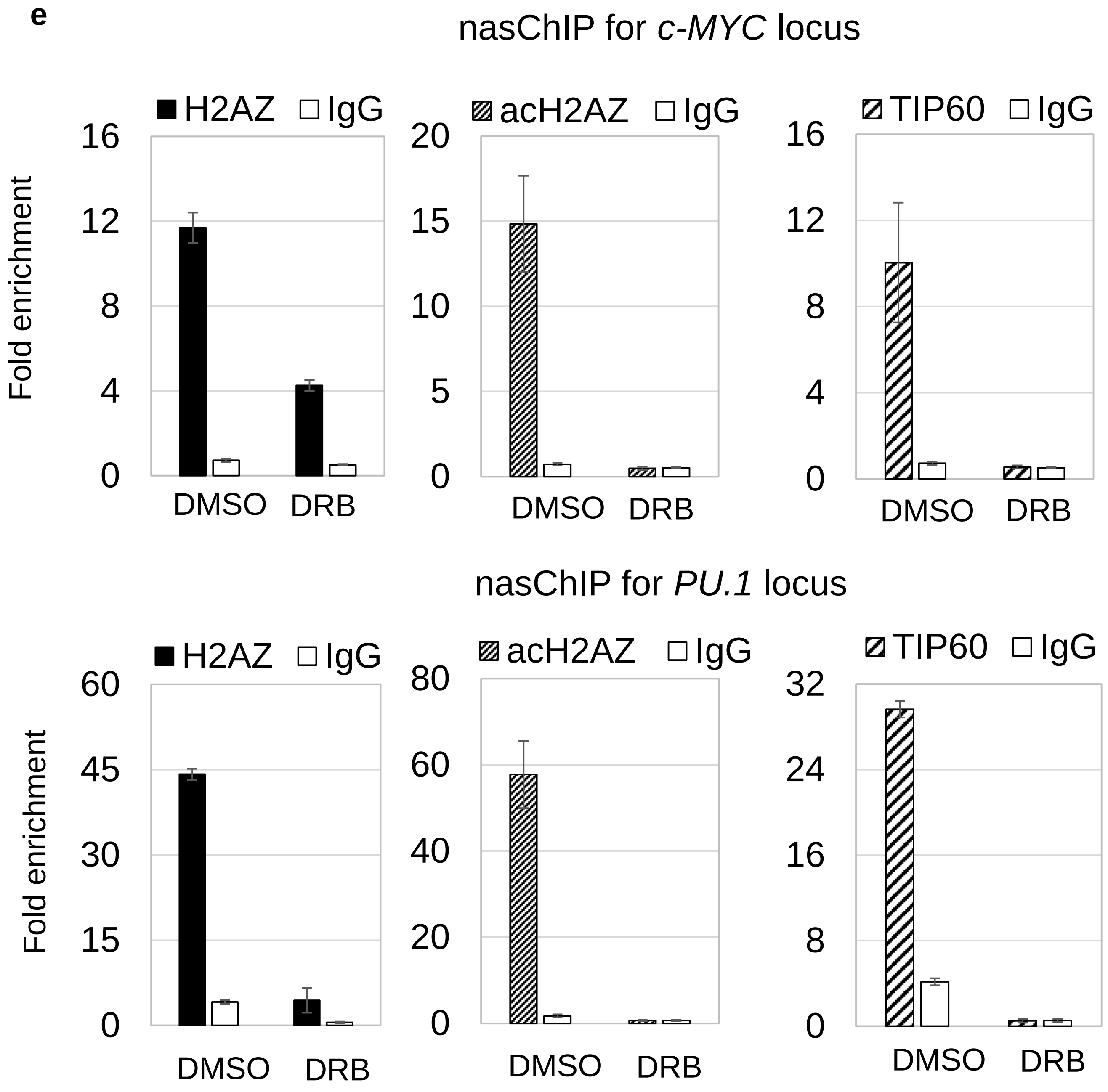


f

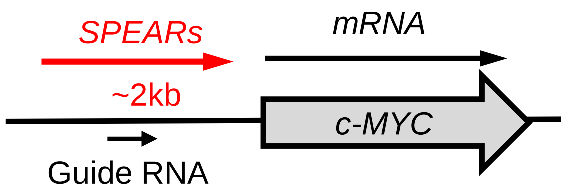


*c-MYC SPEAR* gRNA design

Distance to TSS

ACTTTCGCAAACCTGAACGCGGG

PAM

403 bp

**Extended Data Fig. 4. Global and targeted downregulation and upregulation of *SPEARs* lead to corresponding decreased and increased occupancy of acH2A.Z at the TSSs of the linked genes**.

**a**, Schematic diagram of the pilot experiment showing synchronization of HL-60 cells by double thymidine block followed by treatment with RNA Polymerase Inhibitors. Upon release from double thymidine block, cells were treated with DMSO; Actinomycin D (ActD; RNA Polymerase I, II and III Inhibitor), 0.8 μM; and 5,6-Dichlorobenzimidazole 1-β-D-ribofuranoside inhibitor (DRB; RNAPII Inhibitor), 200 μM. Cells were also spiked with analogs EU and/or EdU for downstream Click-iT conversion. In pilot experiments cells were collected at different time points and total proteins were subjected to Western blot analyses. After 2 hours cells were treated with Ficoll: crosslinked chromatin and RNA were then collected. **b**, Shown are the Western blot analyses with proteins isolated after 2 hours treatment. These experiment revealed that overall global contents of H2A.Z/acH2A.Z and TIP60 were not affected by drug treatment. **c**, Individual *SPEARs* inhibition upon DRB and ActD treatment. qRT-PCR quantitations of the effects of the DRB and ActD on *c-MYC* and *MYB SPEARs*. **d**, Quantitative ChIP-PCR for several amplicons covering the *c-MYC* locus after DRB/ActD treatment. **e**, nasChIP results for the *c-MYC* and *PU.1* loci: DRB-induced downregulation of the *c-MYC* and *PU.1 SPEARs* leads to decreased occupancy of H2A.Z, acH2A.Z and TIP60 at the TSSs of the *c-MYC* and *PU.1* genes. HL-60 cells were released into S Phase and treated with DRB for 2 hours. The medium was supplemented with the EdU DNA analog to enable click collection of nascent DNAs. Chromatin was collected to perform ChIP assays with antibodies to H2A.Z, acH2A.Z, TIP60 and IgG. Nascent DNAs were isolated from the immunoprecipitated chromatin (Experimental details are available in STAR Methods and Figure S1 A) and analyzed by qPCR (bars indicate mean ±s.d.). **f**, Schematic diagram showing the positions of the Guide RNAs for the upregulation of the *cMYC SPEARs* using the CRISPR/dCas9-VP64 gene activation system.

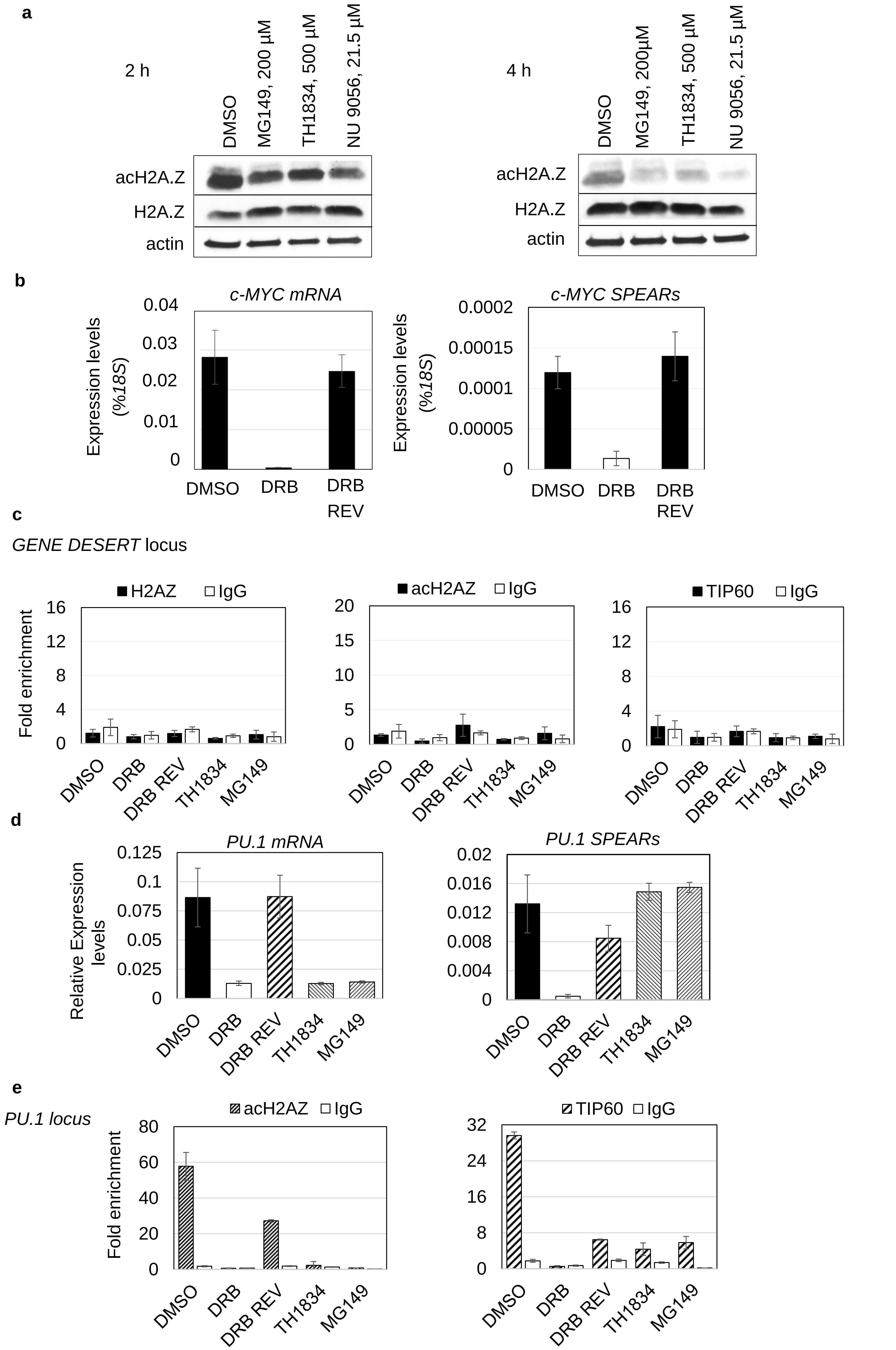


Ebralidze *et.* Extended Data Fig. 5_part 1

**f**

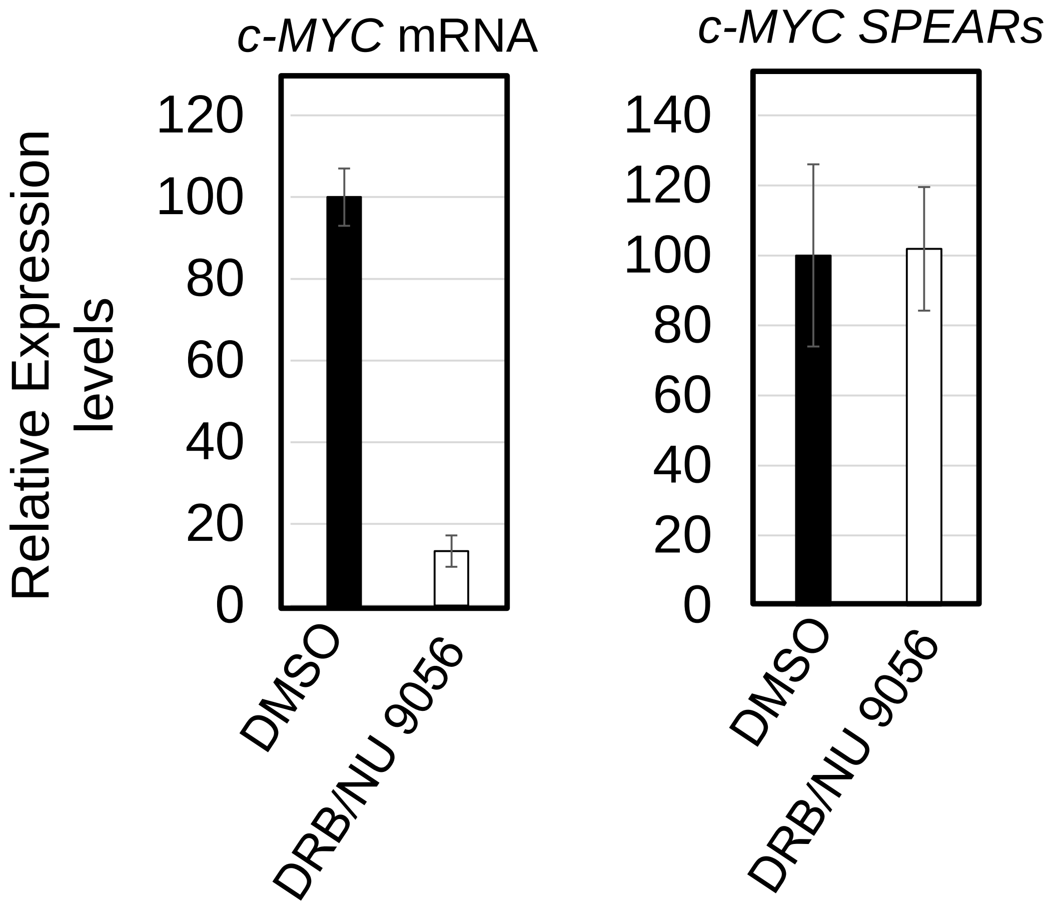
*c-MYC* mRNA
*c-MYC SPEARs*
Relative Expression levels
120
100
80
60
40
20
0
140
DMSO
DRB/NU 9056

**g**

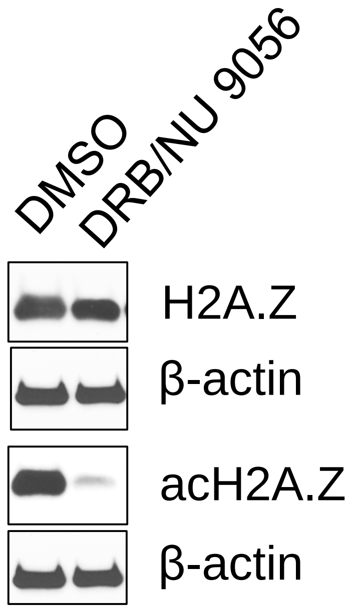
DMSO
DRB/NU 9056
H2A.Z
β-actin
acH2A.Z
β-actin

**h**

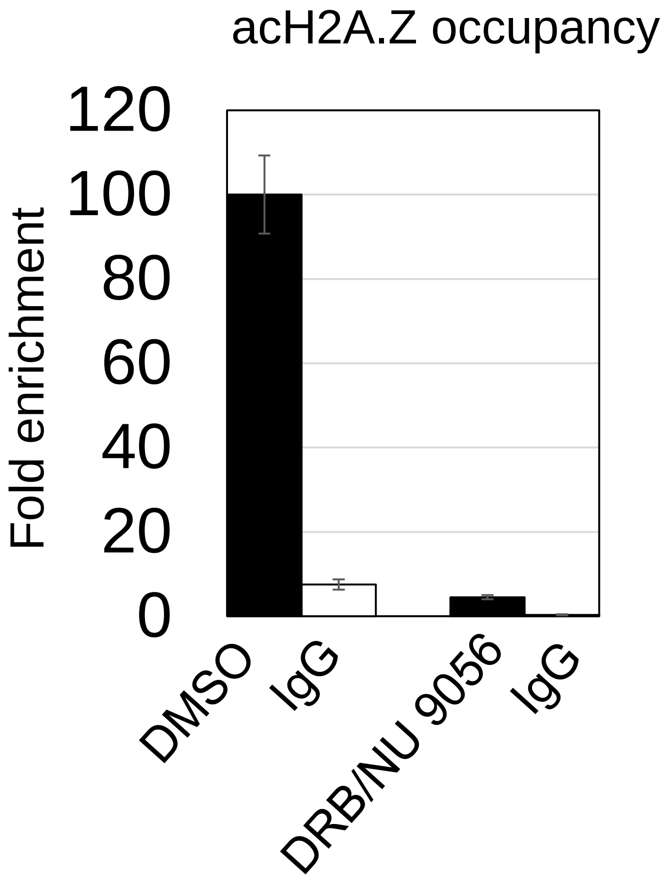
acH2A.Z occupancy
Fold enrichment
120
100
80
60
40
20
0
DMSO
IgG
DRB/NU 9056
IgG

**Extended Data Fig. 5. *SPEARs* regulate the expression of the linked mRNA via TIP60/acH2AZ recruitment/deposition.**

**a**, Upon release from double thymidine block, cells were treated with DMSO; MG149, 200 μM; and TH1834, 250 500 μM; and NU 9056, 21.5 μM. Cells were collected at different time points and total proteins were subjected to Western blot analyses after 2 and 4 hours treatment. **b**, Downregulation of *c-MYC* mRNA*s* and *c-MYC SPEARs* (after DRB treatment), "DRB" bars (middle and right panels) and restoration of *c-MYC* mRNAs and *c-MYC SPEARs* (after DRB reversal), "DRB REV" bars. qRT–PCR and strand-specific qRT-PCR; bars indicate mean +/-s.d. (n=2). **c**, Nascent ChIP-qPCRs for a *GENE DESERT* locus showing no changes in levels of enrichment for H2A.Z, acH2A.Z and TIP60. qPCR, bars indicate mean +/-s.d. (n=2). **d**, Different response of total *PU.1 SPEARs* and mRNA to TIP60/HAT inhibitors: *PU.1* mRNAs are downregulated while *PU.1 SPEARs* are not affected. qRT–PCR and strand-specific qRT-PCR; bars indicate mean +/-s.d. **e**, Nascent ChIP-qPCRs for the *PU.1* locus demonstrating: (i) Downregulation of *PU.1 SPEARs* (after DRB treatment) leads to the loss of acH2A.Z and TIP60 enrichment, "DRB" bars (middle and right panels); (ii) Restoration of of *PU.1 SPEARs* (after DRB reversal) leads to re-occurrence of the acH2A.Z and TIP60 enrichment, "DRB REV" bars (middle and right panels); (iii) Inhibition of TIP60/HAT activity (after MG149 and TH1834 treatments) is preventing the restoration of the acH2A.Z enrichment without affecting TIP60 enrichment, TH1834 and MG149 bars (middle and right panels); qPCR, bars indicate mean +/-s.d. (n=2). **f**, Response of total *c-MYC* mRNA and *SPEARs* after release from synchrony to DRB treatment followed by treatment with the TIP60/HAT inhibitor NU 9056. **g**, Western blot analyses of proteins isolated after consecutive DRB and NU 9056 treatments, showing that the global content of H2A.Z is not affected by DRB and HAT-inhibitor treatment, in contrast to acH2A.Z which is highly depleted, i.e. the modification is lacking.

**h**, ChIP-qPCRs for the *c-MYC* locus using antibodies to acH2A.Z following DRB treatment and HAT inhibition with NU 9056; (amplicon located 652 nt to 345 nt upstream of the *c-MYC* TSS).

**a**

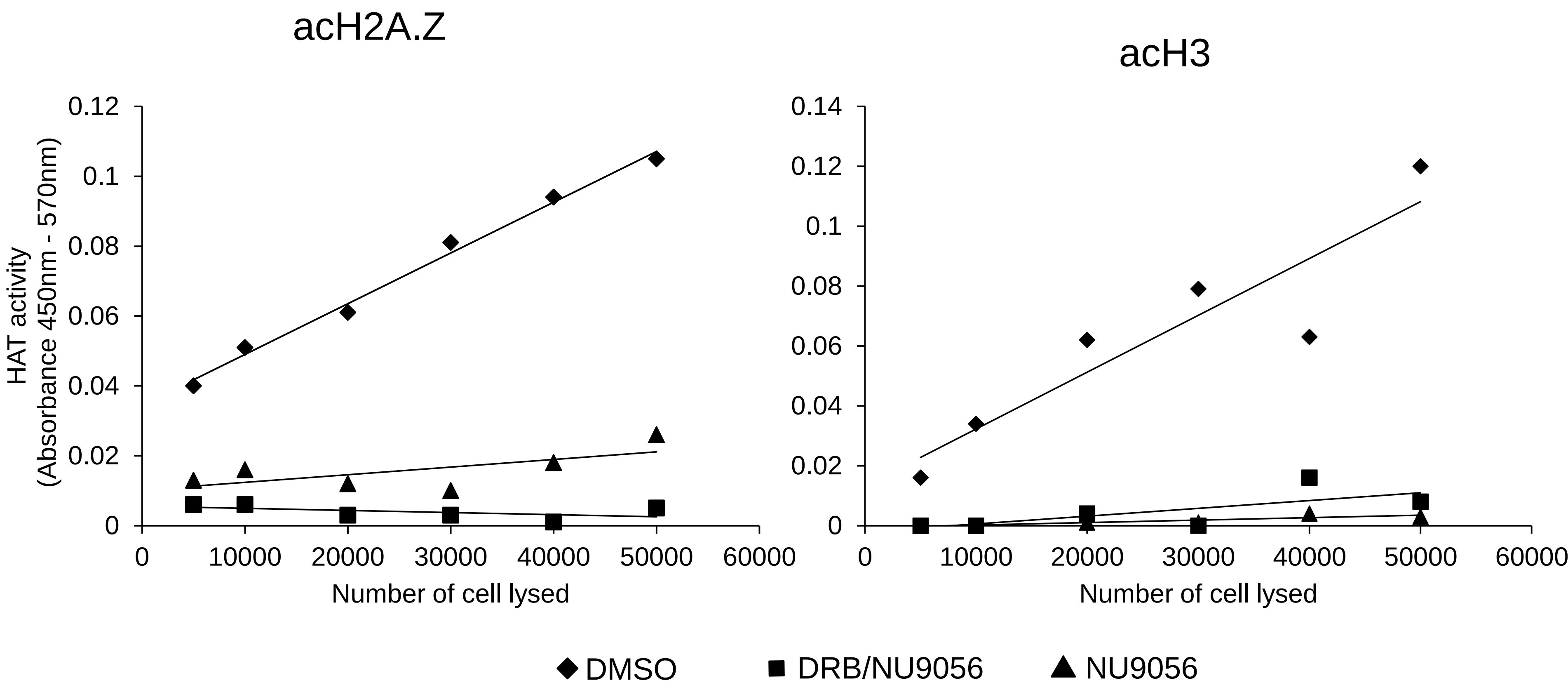


**b**

```
      Ac  Ac   Ac
Ac—AGGKAGKDSGKAKTKAVSRSQRAGLQFP

   VGRIHRHLKSRTTSHGRVGATAAVYSAAIL

   EYLTAEVLELAGNASKDLKVKRITPRHLQL

   AIRGDEELDSLIKATIAGGGVIPHIHKSLIG

   KKGQQKTV
```

**c**

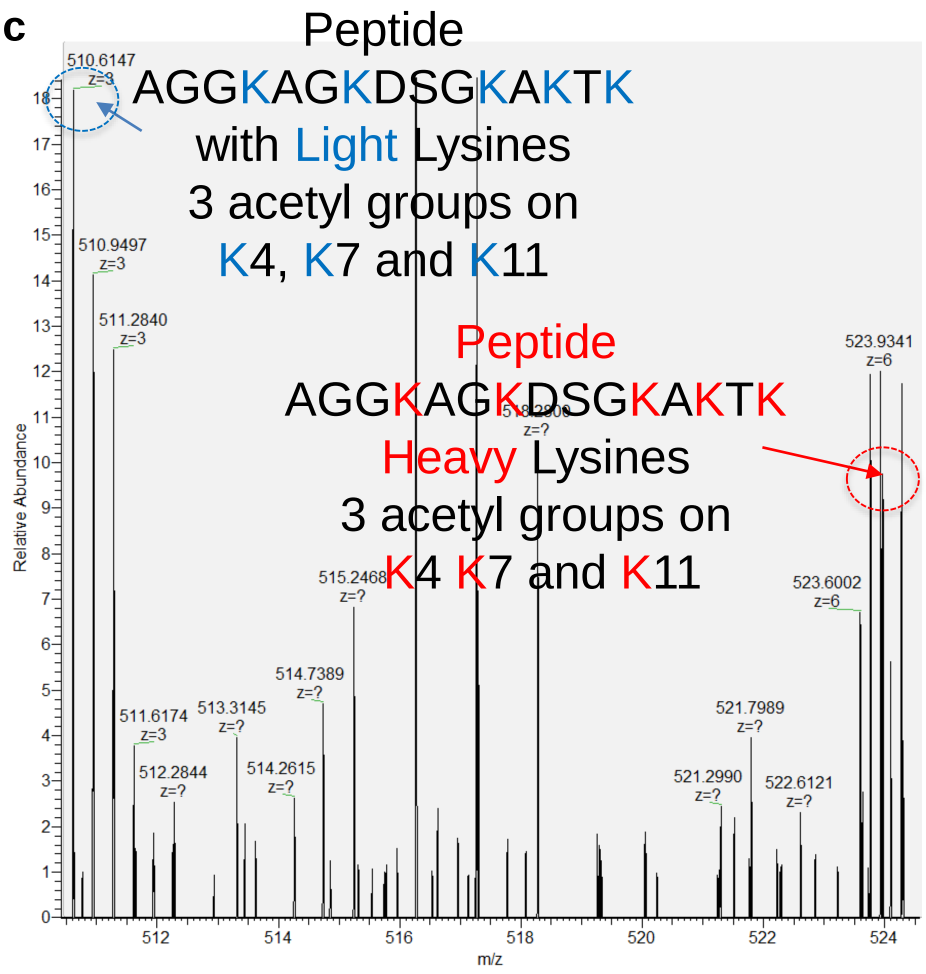


Sample “HATs Inhibitor NU 9056 4 hours”

Ebralidze *et.* Extended Data Fig. 6

**Extended Data Fig. 6. *SPEARs* regulate recycling of the epigenetic mark - acH2AZ**.

**a**, HAT activity assays: 2 and 4 hours of NU 9056 treatment leads to the significant inhibition of global histone acetylation. **b**, Sequence of the H2A.Z. Peptides detectable after Chymotrypsin digestion are shown in red. Shown also position of the acetyl groups in N-terminal peptide. **c**, Example of the detected peptides from sample "HAT inhibitor (NU 9056)-treated sample (4 hours)" after Chymotrypsin digestion of acetylated histone H2A.Z. Light Lysines are marked in blue. Similar peptide was detected in Mock (DMSO)-treated sample (Figure 5 F, right panel).
