## Supplementary material for "Formation and recycling of an active epigenetic mark mediated by cell cycle-specific RNAs": Methods

### Online Methods

#### KEY RESOURCES

##### Mammalian Cell Culture

All cell lines were obtained from ATCC and grown according to the manufacturer's instructions in the absence of antibiotics. The human leukemia cell line HL-60 was grown in RPMI with 10% FBS. The human embryonic kidney cell line HEK293 was cultured in DMEM with 10% FBS.

##### Antibodies

Rabbit polyclonal to human Histone H2A.Z - ChIP Grade (1:2,000; Abcam ab4174), Sheep polyclonal to acetylated human H2A.Z (Ac K4+K7+K11) (1:2,000; Abcam ab18262), Rabbit polyclonal to human TIP60 antibody (generous gift from Bruno Amati).

#### CONTACT FOR REAGENT AND RESOURCE SHARING

Alexander Ebralidze, Annalisa Di Ruscio and Daniel G. Tenen

.

#### METHOD DETAILS

##### RNA isolation

**Total RNA** isolation was carried out as described [1]. All RNA samples used in this study were treated with DNase I (10 U of DNase I per 3 μg of total RNA; 37$^{o}$C for 1 h; in the presence of RNase inhibitor). After DNase I treatment, RNA samples were extracted with acidic phenol (pH

4.3) to eliminate any remaining traces of DNA. cDNA syntheses were performed with Random Primers (Invitrogen) or gene-specific primers with Transcriptor Reverse Transcriptase (Roche Applied Science) according to the manufacturer's recommendation. cDNA was purified with a High Pure PCR Product Purification Kit (Roche Applied Science).

**qRT-PCR**

Sybr green reactions were performed using iQ Sybr Green Supermix (Biorad, Hercules, CA) using the following parameters: 95°C (10 min), 40 cycles of 95°C (15s) and 60°C (1min) 72°C (1min). TaqMan analysis was performed using iTaq Universal Probes One-Step RT-qPCR kit (Biorad, Hercules, CA) at the following conditions: 50°C (10 min.), 95°C (2 min.) and then 40 cycles of 95°C (15 sec.) and 60°C (60 sec.).

**Primers for TaqMan real time PCR:** Human MYC mRNA: ABI Cat. # Hs00153408_m1; Human MYB mRNA: ABI Cat. # Hs00920556_m1; *c-MYC SPEARs*: PCR primers: Forward: 5'- ACA CAT CTC AGG GCT AAA CAG -3'; Probe: 5'- ATA CCT TCC ACC CAG ACT GAG TCC C -3', Reverse: 5'- TGC ACA GCT ATC TGG ATT GG -3'.

**Primers used for strand-specific real-time RT PCR (Sybr):** Reverse Transcriptase primer for *c-MYC SPEARs*: PCR primers: Forward: 5'- ACA GGC AGA CAC ATC TCA GGG CTA -3'; Reverse: 5'- ATA GGG AGG AAT GAT AGA GGC ATA -3'.  Reverse Transcriptase primer for *PU.1 SPEARs*: 5'- GGC TTT TGC TCT AAC CCA AC -3'; PCR primers: Forward: 5'- ACT ATG CTG AAG ACC CTA CAC -3'; Reverse: 5'- GCT CTA ACC CAA CAA ATG CC -3'.

**Nascent RNA/DNA capture** was performed using Click-iT® Nascent RNA Capture Kit (ThermoFisher) according to the manufacturer's instructions with minor modifications. Briefly, 1. **Labeling the cells with EU/EdU**. 200 mM EU or 30 mM EdU stock solutions were added to the cells, to a final concentration 0.5 mM or 30 μM, respectively. 2. **Incubation**. The cells were incubated for 1 or 2 h. 3. **RNA/DNA isolation**. The cells were harvested and the RNA/DNA were isolated and dissolved in 14 μL of $H_2O$. 4. **Biotinylation of RNA/DNA by Click reaction**. Click-iT® reaction cocktail (50 μL per reaction) was prepared accordingly to manufacturer's instructions: a mixture containing 1x Click-iT EU buffer; 2 mM $CuSO_4$; 1 mM Biotin azide; 13.25 μL of the isolated RNA; 10 mM Click-iT® reaction buffer Additive 1; 12 mM Click-iT® reaction buffer additive 2 was prepared. After adding each component, the reaction cocktail was gently mixed by vortexing. The addition of the Click-iT® reaction buffer Additive 1 stock initiates the click reaction between the EU-RNA/EdU-DNA and biotin azide. Subsequently, the Click-iT® reaction buffer Additive 2 is added and incubated for 30 min with gentle vortexing. 5. **RNA/DNA precipitation**. 1 μL of UltraPure™ Glycogen, 55 μL of 7 M ammonium acetate, and 750 μL of chilled 100% ethanol were added to the click reaction, incubated at –70°C for at least 30 min and after centrifugation the pellet was dissolved in 125 μL of $H_2O$. 6. **Binding biotinylated RNA/DNA to Dynabeads® MyOne™ Streptavidin T1 magnetic beads (ThermoFisher)**. The RNA/DNA binding reaction mixture included: 125 μL 2xClick-iT® RNA binding buffer; 2 μL Ribonuclease Inhibitor or 2 μL of water for DNA; 125 μL of the isolated biotinylated RNA/DNA. The RNA binding reaction mixture was heated at 68–70°C for 5 min and 50 μL of bead suspension added into the heated RNA binding reaction mixture. The tube containing the RNA/DNA binding reaction was incubated at r.t. for 30 min while gently vortexing to prevent the beads from settling. The beads were immobilized using the magnet and washed 5 times with 500 µL of Click-iT® reaction wash buffer 1 and 5 times with 500 µL of Click-iT® reaction wash buffer 2. Finally, the beads were resuspended in 50 µL of Click-iT® reaction wash buffer 2 and the captured RNA immediately reverse transcribed to cDNA. The captured DNA was released into 50 µL of boiling water and used in qPCR analyses.

**Primer extension and 5'/3' RACE** cDNAs from the HL-60 cell line were synthesized as described above and run in urea-PAGE [2]. **5'/3' RACE** was performed using the Exact START™ Eukaryotic mRNA 5'- & 3'-RACE Kit (Cambio Ltd, UK) according to the manufacturer's instructions.

**Double Thymidine block (early S-phase block)** was carried out as described [3]. Briefly, HL-60 cells were grown overnight to 70-80% confluence, washed twice with 1xPBS and cultured in DMEM (10% FCS) + 2.5 mM Thymidine for 18 hrs (first block). Thymidine was washed out with 1xPBS and cells were grown in DMEM (10% FCS). After 8 h cells were cultured in the presence of thymidine for 18 h (second block) and then released as described [3]. Synchrony was monitored by flow cytometry analysis of propidium iodide-stained cells using a LSRII flow cytometer (BD Biosciences) at the Harvard Stem Cell Institute/Beth Israel Deaconess Center flow cytometry facility.

**DRB and Actinomycin D treatments** were carried out as described [3]. Briefly, after release from double thymidine block, HL-60 cells were treated with 100 µM of 5,6-Dichlorobenzimidazole 1-β–D-ribofuranoside (DRB, [4]; Sigma Aldrich, or 0.8 µM of Actinomycin D (Sigma Aldrich) as indicated at each time point.

**Down-regulation of *c-MYC SPEARs***

27 siRNAs (Small interfering RNAs) targeting the *c-MYC SPEARs* were designed and synthesized by siTOOLs Biotech and designated as *c-MYC SPEARs* siPool (*siMYC*) [5]: the sequences are shown in the following Table A. The *siMYC* were dissolved in nuclease-free water and used to transfect HL-60 cells using Amaxa Cell Line Nucleofector Kit V, Program T-019 (Nucleofector® II Device; Lonza) according to the manufacturer's instructions. Briefly, $2x10^6$ cells/reaction were cultured in RPMI medium +10% FBS. The cells were collected by centrifugation. 1 μL si*SPEARs* and Negative Control siPool (siControl) was mixed with 4 μL nuclease-free water per reaction (siPool solution). Cells were resuspended in 100 μL Nucleofector® Solution per reaction at room temperature and combined with the siPool solution. The final concentration of si*SPEARs* and siControl was 100 nM. The cells/*siMYC* and cells/siControl suspensions were transferred into certified cuvettes and taken through Nucleofector® Program T-019 (Nucleofector® II Device) in triplicates. 500 μL of the pre-incubated culture medium was immediately added to the cuvette and the samples were gently transferred into the wells of 6-well plate containing 500 μL of the pre-incubated culture medium. The samples were cultured for 24 h, the electroporation was repeated and then cultured for a further 24 h. Live cells were harvested with Ficoll-Paque PLUS medium (GE Healthcare, # 17144003), and RNA/chromatin extracted.

Small interfering RNAs Sequences are indicated in **Supplementary Table A**.

**Up-regulation of *c-MYC SPEARs***

Guide RNA (gRNA) targeting the *c-MYC SPEARs* was designed on the Genetic Perturbation Platform (https://portals.broadinstitute.org/gpp/public/analysis-tools/sgrna-design). A lentiviral vector (pLenti-U6-sgRNA-PGK-Neo) encoding the designed gRNA was synthesized by Applied Biological Materials Inc. pMD2.G, pCMV delta R8.2, and the lentivector were transfected into 10 million 293T using TransIT-LT1 reagent (Mirus). Supernatants were harvested at 48 and 72 hrs after transfection. Virus was concentrated 100 times by Lenti-X Concentrator (Takara) after filtering through a 0.45 μm syringe filter, and resuspended in DMEM and stored at –80$^{0}$C. dCas9-VP64 (Addgene, plasmid #61425) lentiviral transduction was performed in the presence of hexadimethrine bromide (final concentration 8 μg/ml) in HEK 293. Blasticidin (10 μg/ml) was added to the cultures 2 days after infection. Stably expressing dCas9-VP64 HEK 293 cells were subsequently transduced with *c-MYC SPEAR* gRNA or scrambled control. G418 (500 μg/ml) was added to the cultures 2 days after infection. Resistant clones were selected and screened for *c-MYC SPEARs* as well as *c-MYC* levels. The gRNA sequence is indicated in Extended Data Fig.4f.

**Histone Acetyltransferases (HAT) Assay**

Cells were lysed in lysate buffer (50 mM Tris, pH8; 10% Glycerol; 0.1 mM EDTA; 1 mM DTT; 10 mM sodium butyrate, 0.1% Triton X-100, protease inhibitor cocktail). Cell lysates were sonicated and supernatants collected by spinning at 5000 g for 5 min at 4°C. 100 μl of 1 ng/μl biotinylated histone peptide (H2A.Z and H3 in this experiment) was added to each well of streptavidin-coated plates (Thermo Fisher, #15121). After incubation for 30 min at room temperature, plates were washed with TBS buffer. Plates were blocked with 200 μl of 3% BSA in TBS for 30 min at 30°C. HAT assay was performed by adding 50 μl of HAT reaction cocktail to each well (50 mM Tris, pH=8; 10% Glycerol; 0.1 mM EDTA; 1 mM DTT; 1 mM Acetyl-CoA; cell lysate), and plates were incubated for 30 min at 30°C. Plates were then stained for 1.5 hrs at room temperature with 100 µl of the following primary antibodies in TBS/3%BSA: H2A.Z acetyl (K4+K7+K11) (1:500; Abcam ab18262), H3 acetyl (1:500, Sigma Aldrich #06-599). 100 µl of secondary horseradish peroxidase (HRP)-conjugated antibodies in TBST/3% BSA were diluted 1:10,000 and added to each well for 30 min at room temperature. 100 µl of a 1:1 mixture of 0.006% $H_2O_2$ and TMB solution (Thermo Scientific, #N301) was added to each well for 30 min in the dark. 50 µl of fresh 1M sulfuric acid was added to each well to stop the HRP reaction. The mixture was transferred to 96 well plates, which were read on a plate reader at a wavelength of 450 nm and 570 nm. 570 nm values were subtracted from 450 nm values to remove any well-to-well variation.

**Ribonucleoprotein (RNP) fractionation and RNP pull-down assay**

Equal numbers of viable cells counted after Ficoll gradient purification, were used for each isolation. Nuclei from $2x10^6$ cells were isolated as described [6]. Briefly, equal amounts of viable cells were washed with ice-cold PBS supplemented with 5 mM vanadyl complex, 1 mM PMSF, and resuspended in ice-cold lysis buffer: 1x Buffer A (10 mM HEPES-NaOH pH 7.6; 25 mM KCl; 0.15 mM spermine; 0.5 mM spermidine; 1 mM EDTA; 2 mM Na butyrate); 1.25 M sucrose; 10% glycerol; 5 mg/mL BSA; 0.5% NP-40; freshly supplemented with protease inhibitors (2 mM leupeptin; 2 mM pepstatin) 100 mM benzamidine; protease inhibitor cocktail (Roche Applied Science, Cat. No. 1836153), 1 tablet/375 µL $H_2O$; 100 mM PMSF; 2 mM vanadyl complex (New England Biolabs) and 20 units/mL RNase inhibitor (RNAguard; Amersham Biosciences). Samples were incubated at $0^{\circ}$C for ~10 min and subjected to several strokes in a Dounce homogenizer. The pelleted nuclei were resuspended in 0.5 ml lysis buffer and diluted with 2.25 mL Dilution Buffer (2.13 mL "Cushion" buffer plus 0.12 mL 0.1 g/mL BSA), freshly supplemented with protease inhibitors and overlaid onto 2 mL "cushions" (200 mL "Cushion" buffer consists of 15 mL dd$H_2O$; 15 mL 20x Buffer A; 30 mL glycerol; 240 mL 2.5 M sucrose; freshly supplemented with protease inhibitors) into one SW 55 Ti tube and centrifuged at 24,400 rpm, for 60 min at 4$^o$C. After washing with PBS/1mM PMSF, nuclei were then resuspended in 1.8 ml of cytoskeletal buffer (CSK-50: 10 mM Pipes, pH 6.8; 300 mM sucrose; 50 mM NaCl; 3 mM $MgCl_2$; 1 mM EGTA; 0.5 mM vanadyl complex; 1 mM PMSF). 20 µL (200 U) DNase I was added and incubation carried out at 37$^o$C for 30 min and then chilled on ice. After addition of 200 µL of 2.5 M $(NH_4)_2SO_4$ (final 250 mM), the nuclei were incubated on a rocking platform 40 min at 4$^o$C and spun down at 2,000 g for 5 min at 4$^o$C. The supernatant fraction was discarded. The RNP-containing pellet was washed twice with ice-cold PBS and resuspended in 2 mL RIP buffer #4 (50 mM HEPES-KOH, pH 7.5, 100 mM NaCl, 1 mM EDTA, 0.5 mM EGTA, 0.1% Na-Deoxycholate, 0.5% N-lauroylsarcosine, 1× protease inhibitors.). After solubilization by sonication, the pellet was chilled on ice and spun down at 5,000 g for 5 min at 4$^o$C. The supernatant – the RNP fraction (~2 ml each) was collected and used in RNP pull-down assay and SDS-PAGE/Western Blotting analysis. RNP pull-down assay was performed using the Pierce$^{TM}$ Magnetic RNA-Protein Pull-Down Kit (Thermo Scientific; Cat. # 20164) according to manufacturers' recommendation (http://bit.ly/3qNxH8L). SDS-PAGE/Western Blotting analysis was performed as follows: 1.2 µL of 20% SDS; 0.8 µL 1 M DTT solution were added to 40 µL of RNP fractions and boiled for at least 1 min. Samples were desalted on a pre-equilibrated G25 column (with 100 mM Tris-HCl, pH 7.4; 10 mM DTT; 0.6% SDS) and analyzed SDS-PAGE/Western Blotting.

**Western Blotting Analysis**

Whole-cell lysates from approximately $0.2x10^6$ cells per sample were separated on 12% SDS-PAGE gels and transferred to a nitrocellulose membrane. Immunoblots were stained overnight at 4°C with the following primary antibodies: H2A.Z (1:2,000; Abcam ab4174 ), H2A.Z acetyl (K4+K7+K11) (1:2,000; Abcam ab18262), or TIP60 (1:1000; generous gift from Bruno Amati). Secondary horseradish peroxidase (HRP)-conjugated antibodies were diluted 1:5,000 and incubated for 1h at room temperature with TBST/5% BSA. Immuno-reactive proteins were detected using the Pierce® ECL system (Thermo Scientific #32106).

**Tandem Mass Spectrometry (LC-MS/MS)**

Whole cell extracts were separated on SDS-PAGE gels which were stained with Coomassie brilliant blue and the protein bands of interest excised. Gel sections were reduced with 55 mM DTT, alkylated with 10 mM iodoacetamide (Sigma-Aldrich), and digested overnight with TPCK modified trypsin/LysC mix or Chymotrypsin, Sequencing Grade (Promega) at pH=8.3. Digestion was stopped with 1% TFA and peptides dried down in a SpeedVac to 10 µL. The peptide mixture was analyzed by positive ion mode LC-MS/MS using a high-resolution hybrid QExactive HF Orbitrap Mass Spectrometer (Thermo Fisher Scientific) via HCD with data-dependent analysis (DDA). Peptides were delivered and separated using an EASY-nLC nanoflow HPLC (Thermo Fisher Scientific) at 300 nL/min using self-packed 15 cm length × 75 µm i.d. C18 fritted microcapillary columns. Solvent gradients were: 90 min from 3% to 38% buffer B (100% acetonitrile) in buffer A: (0.9% acetonitrile/0.1% formic acid/99.0% water). The raw files were processed with MaxQuant version 1.5.2.8 [7] with preset standard settings at a multiplicity of 1. Carbamidomethylation was set as a fixed modification while methionine oxidation and protein N-acetylation were considered as variable modifications. Search results were filtered with a false discovery rate of 0.01. Reverse hits and only by site identifications as well as potential contaminants were removed. MS data will be deposited to the ProteomeXChange Consortium via PRIDE upon acceptance of the manuscript. Private partial submission with MASSIVE:

https://massive.ucsd.edu/ProteoSAFe/dataset.jsp?task=38a5eae58dd247f5b3c42f2be181e692

Password: Alex_Acetyl

**Nuclear Chromatin (ChIP) and RNA immunoprecipitation (nRIP)**

**ChIP** was performed as follows. Cells were crosslinked with 1% formaldehyde for 10 min at room temperature. Pellets of $1x10^6$ cells were used for immunoprecipitation as previously described [8] and lysed for 10 min on ice and chromatin fragmented using a Branson 250 digital sonicator. Each ChIP was performed with 4ug of antibody, incubated overnight at 4°C. A 50/50 slurry of protein A and protein G Dynabeads was used to capture enriched chromatin, which was then washed before reverse-crosslinking and proteinase K digestion at 65C. AMPure XP beads were used to clean up and isolate ChIP DNA for subsequent library construction. The following antibodies were used for ChIP: H2A.Z (Abcam ab4174, lot GR3176820-1), acH2A.Z (Abcam ab18262, lot GR306397-1), TIP60 antibody[9] (generous gift from Bruno Amati); and IgG (Abcam ab171870). Fold enrichment was calculated using the formula $2^{(-\Delta\Delta Ct(ChIP/non\text{-}immune\ serum))}$. Primer sets used for ChIP are listed in **Supplementary Table B**.

**nRIP** was performed as described [6] with some modifications. Crosslinked **nuclei** were collected as follows: 1. $60x10^6$ HL-60 cells were crosslinked with 1% freshly made formaldehyde solution:

50 mM HEPES-KOH; 100 mM NaCl; 1 mM EDTA; 0.5 mM EGTA; 11% formaldehyde) for 10 min at room temperature. 2. Crosslinking was stopped by adding 1/10$^{th}$ volume of 2.66 M Glycine, kept for 5 min at room temperature and 10 min on ice. 3. Cell pellets were washed twice with ice-cold PBS (freshly supplemented with 1 mM PMSF). 4. Cell pellets were resuspended in cell lysis buffer (volume = 4 mL): 1x Buffer (10 mM Tris pH 7.4; 10 mM NaCl; 0.5% NP-40), freshly supplemented with protease inhibitors (protease inhibitors cocktail: Roche Applied Science, Cat. No. 1836153, 1 tablet/375 μL $H_2O$; add as x100), 0.1 mM PMSF, and 0.2 mM vanadyl complex (NEB). 5. Cells were incubated at 0°C for 10-15 minutes and homogenized in a Dounce (10 strokes pestle A and 40 strokes pestle B). 6. Nuclei were recovered by centrifugation at 2,000 rpm for 10 min at 4°C. 7. Nuclei were resuspended in 3 ml of 1x Resuspension Buffer (50 mM HEPES-NaOH, pH 7.4; 10 mM $MgCl_2$) supplemented with 0.1 mM PMSF and 0.2 mM vanadyl complex. 8. DNase I treatment (250 U/ml) was performed for 30 min at 37°C, and EDTA (final concentration 20 mM) added to stop the reaction. 9. Resuspended Nuclei were sonicated once for 20s (1 pulse every 3 seconds) at 30% amplitude (Branson Digital Sonifer, Danbury, CT).

**Immunoprecipitation** was performed as follows: 1. Before preclearing, the sample was adjusted to 1% Triton X-100; 0.1% sodium deoxycholate; 0.01% SDS; 140 mM NaCl; Protease inhibitors; 0.2 mM vanadyl complex; 0.1 mM PMSF. 2. Preclearing step: ~ 50 μL magnetic beads (Protein A or G Magnetic Beads; #S1425S or #S1430S NEB) were added to the sample and incubation was carried out for 1 h on a rocking platform at 4°C. 3. Beads were removed in the magnetic field. 4. The sample was then divided into five aliquots: (i-iii) antibody of interest: (i) H2A.Z antibody (ab4174); (ii) acH2A.Z antibody (ab18262); (iii) TIP60 antibody [9] (generous gift from Bruno Amati); (iv) preimmune serum: IgG (ab171870); (v) no antibody, no serum (input). 5. 5 μg antibody or preimmune serum was added to the respective aliquot and incubation performed on a rocking platform overnight at 4°C. Input was stored at -20 °C after addition of SDS to 2% final concentration. Day II. 6. 200 μL of Protein A coated super-paramagnetic beads (enough to bind 8 μg IgG) were added to the samples and incubated on a rocking platform for 1 h at 4°C. 7. Six washes of beads in the magnetic field were made with immunoprecipitation buffer (150 mM NaCl; 10 mM Tris-HCl, pH 7.4; 1 mM EDTA; 1 mM EGTA pH 8.0; 1% Triton X-100; 0.5% NP-40 freshly supplemented with 0.2 mM vanadyl complex and 0.2 mM PMSF) in a magnetic field. 8. Proteinase K treatment to release DNA/RNA into solution and to reverse the crosslinking was performed in 200 μL of: 100 mM Tris-HCl, pH 7.4; 0.5% SDS for the immunoprecipitated samples and in parallel for the input using 500 μg/mL of Proteinase K at 56°C overnight. 9. Day III. Beads were removed in the magnetic field. 10. Phenol (pH 4.3) extraction was performed after addition of NaCl (0.2 M final concentration). 11. Ethanol precipitation (in the presence of glycogen); 3 h at -20°C. 12. The pellet was dissolved in 180 μL $H_2O$, heated at 72 °C for 2 min, and immediately chilled on ice.13. Samples were treated with DNase I (250 U/ml) in the presence of RNase inhibitor at 300 U/ml in x1 buffer # 2 (NEB) at 37°C for 30 min. 14. Phenol (pH 4.3) extraction and EtOH precipitation were repeated. 15. The RNA pellet was dissolved in 50 μL $H_2O$.

**RNA electrophoretic gel mobility shift assays (REMSAs)**

RNA oligonucleotides (15 pmol) were end-labeled with [$\gamma$-$^{32}$P] ATP (Perkin Elmer) and T4 polynucleotide kinase (New England Biolabs). Reactions were incubated at 37°C for 1 h and then passed through G-25 spin columns (GE Healthcare) according to the manufacturer's instructions to remove unincorporated radioactivity. Labeled samples were gel-purified on 10% polyacrylamide gels. Binding reactions were carried out in 10 μL volumes in the following buffer: 5 mM Tris pH 7.4, 5 mM $MgCl_2$, 1 mM DTT, 3% v/v glycerol, 100 mM NaCl. 5μg of full length purified H2A.Z (Abcam) and TIP60 (SignalChem) proteins were incubated with 1.1 nM of $^{32}$P-labeled single-stranded (ss) RNAs. All reactions were assembled on ice and then incubated at room temperature for 30 min. Samples were loaded onto 6% native polyacrylamide gels (0.5xTBE) at 4 °C for 3h at 140 V. Various concentrations (1μM-10mM) of H2A.Z and K7 acetylated H2A.Z peptides (AnaSpec) were incubated with a fixed amount of probe in 50 mM Tris-HCl pH 7.5, 100 mM NaCl, 0.4 mM EDTA, and 40 U/ml RNasin,1 mM DTT, 50% glycerol in a final volume of 10 μL. Thereafter, 1 μl glutaraldehyde (0.2% final concentration) was added into the mixture and incubated at room temperature for 15 minutes. Samples were loaded onto 10% native polyacrylamide gels (0.5xTBE) at 4 °C for 3h at 170 V. All gels were dried upon fixation with methanol and acetic acid and exposed to X-ray film. RNA oligonucleotides are listed in **Supplementary Table C.**

#### QUANTIFICATION AND STATISTICAL ANALYSIS

##### RIP-Sequencing Analyses

Immunoprecipitated RNA were processed for sequencing as described by Di Ruscio *et al*.[3] with some modifications. RNA samples were depleted of ribosomal RNA with Ribo-ZeroTM Magnetic Gold Kit (cat. # MRZG126 Epicentre). Double stranded cDNA libraries were constructed using ScriptSeq™ v2 RNA-Seq Library Preparation Kit (cat. # SSV21106 Epicentre). The libraries were subjected to final size-selection in 3% agarose gels. 250-500 bp fragments were excised and recovered using the Qiaquick Gel Extraction Kit (Qiagen). Raw fastq files had optical duplicates removed using clumpify from BBMap (https://sourceforge.net/projects/bbmap/) with the flags "dedupe spany addcount". Next, Adaptor Trimming was performed using BBDuk (from BBMap) with the flags "ref=/.../bbmap/resources/adapters.fa ktrim=l hdist=2". Next, reads were trimmed using trimmomatic [10] with the flags "LEADING:24 SLIDINGWINDOW:4:24 TRAILING:24 MINLEN:20". After cleanup, reads were aligned to hg38 using STAR v2.7.5a [11] with the flags " --readFilesCommand zcat --outSAMtype BAM SortedByCoordinate". Bam file sorting and indexing was performed using samtools [12]. Bam files were split by strand using bamtools' [13] filter command. Bam coverage maps were generated using bamCoverage from deeptools [14] using default parameters with the exception of bin size being set to 25bp. These coverage maps have been uploaded to GEO. Peak calling was performed on all samples using MACS2 [15]( HOMER [16]( Genrich (https://github.com/jsh58/Genrich) and SICER2 (https://github.com/zanglab/SICER2) . The table below gives peak caller settings for each RIP sample that differed from the default parameters. A blank cell represents peak calling with default parameters.

| Peak Caller | acH2Az (Forward) | TIP60 (Forward) | H2Az (Forward) |
|---|---|---|---|
| MACS2 | --slocal 3700<br>--broad | --slocal 5000<br>--broad | --slocal 3000<br>--broad |
| HOMER | -style histone<br>-region<br>-size 150<br>-minDist 370 | -style histone<br>-region<br>-size 150<br>-minDist 370 | -style histone<br>-region<br>-size 150<br>-minDist 370 |
| Genrich | | | |
| SICER2 | | | |

| Peak Caller | acH2Az (Reverse) | TIP60 (Reverse) | H2Az (Reverse) |
|---|---|---|---|
| MACS2 | --slocal 3200<br>--broad | --slocal 5000<br>--broad | --slocal 2800<br>--broad |
| HOMER | -style histone<br>-region | -style histone<br>-region | -style histone<br>-region |

| | -size 150<br>-minDist 370 | -size 150<br>-minDist 370 | -size 150<br>-minDist 370 |
|---|---|---|---|
| Genrich | | | |
| SICER2 | | | |

Once called, peaks were merged using HOMERs mergePeaks command with the flag "-d 500." Average fold change of each peak over input was calculated using a custom python script and considering only uniquely mapped reads.  Peaks were also annotated using HOMER's annotatePeaks command using the hg38 genome. For peak filtering to select for peaks within - 2kb - +1kb of the TSS, the reported "Distance to TSS" value was used and this resulted in the list of "putative SPEARs". Genome coverage plots were generated using ngsplot [17] with the flags "-G hg38 -R tss -L 2000 -RB 0.05" and tailored y-axis values per plot.

For generating the SPEAR coverage regions, normalized bam coverage maps were generated with bamCoverage using the flags "--outFileFormat bedgraph --normalizeUsing RPGC --binSize 25 --effectiveGenomeSize 2864785220". The generated bedgraphs were then imported into R v4.0.3 (https://www.R-project.org/ and figures generated using the package Sushi (https://github.com/dphansti/Sushi).

**RNA Sequencing**

RNA was extracted with TRI Reagent® (MRC). RNA samples were treated with DNase I: 10 U of DNase I (Roche) per 3 μg of total RNA; 37ºC for 1 hr in the presence of RNase A inhibitor. RNAs were depleted of ribosomal RNA with Ribo-ZeroTM Magnetic Gold Kit (cat. # MRZG126 Epicentre). Double stranded cDNA libraries were constructed using ScriptSeq™ v2 RNA-Seq Library Preparation Kit (cat. # SSV21106 Epicentre). The libraries were subjected to final size-selection in 3% agarose gel. 250-500 bp fragments were excised and recovered using the Qiaquick Gel Extraction Kit (Qiagen). Libraries were sequenced on Hi-Seq-2000 Illumina.

Raw fastq files had optical duplicates removed using clumpify from BBMap (https://sourceforge.net/projects/bbmap/) with the flags "dedupe spany addcount". Adaptor trimming was performed using BBDuk (from BBMap) with the flags "ref=/.../bbmap/resources/adapters.fa ktrim=l hdist=2". Reads were then trimmed using trimmomatic (REF) with the flags "LEADING: 28 SLIDINGWINDOW: 4:26 TRAILING: 28 MINLEN: 20". After cleanup, reads were aligned to hg38 using STAR v2.7.5a [11] with the flags " --readFilesCommand zcat --outSAMtype BAM SortedByCoordinate". Bam file sorting and indexing was performed using samtools. Bam coverage maps were generated using bamCoverage [12] using default parameters. Post-alignment QC was performed using RSeqQC [18] to assess gene body coverage. Transcript quantification was performed using featureCounts v2.0.0 [19] using the flags "-g gene_id --extraAttributes gene_name -M -O -d 20."

To generate the list of "expressed" genes, raw counts were read into R v4.0.3 and counts-per-million (cpm) values determined for all genes. CPM values less than 1 (which corresponded to raw read counts of ~6-8) were counted as "non-expressed" genes and had their cpm set to 0 for further analysis. This was done to remove lowly expressed genes from the dataset which would confound further analysis. This list of "expressed" genes was then merged with the putative SPEARs listing from the RIP analysis merge based on Ensembl_ID. The scatter plot presented showing Nuclear vs Nuclear Clicked expression was generated using the R package ggplot2 (https://ggplot2.tidyverse.org).

**ChIP- Sequencing Analyses**

ChIP library construction was performed as described [8] and paired-end sequenced on the NextSeq500 platform, at a reading length of 36 nucleotides. For the DMSO/DRB/ActD and siMYC ChIP-Seq analyses, raw fastq files had optical duplicates removed using clumpify from BBMap (https://sourceforge.net/projects/bbmap/) with the flags "dedupe spany addcount." Adaptor trimming was performed using BBDuk (from BBMap) with the flags "ref=/.../bbmap/resources/adapters.fa ktrim=l hdist=2" and reads were trimmed using trimmomatic [10] with the flags "LEADING: 20 SLIDINGWINDOW: 4:20 TRAILING: 20 MINLEN: 20". After cleanup, reads were aligned to hg38 using bwa mem v0.7.17-r1188 [20] with default settings. Bam file sorting and indexing was performed using samtools [20]. Bam coverage maps were generated with bamCoverage (REF) using default parameters with the exception of the bin size being set to 25bp. These coverage maps have been uploaded to GEO. Peak calling was performed on all samples using MACS2 [15] HOMER [16], Genrich (https://github.com/jsh58/Genrich) and GEM(https://groups.csail.mit.edu/cgs/gem). MACS2 was run with default parameters, HOMER was run with the flags "-region -size 150 -minDist 370," Genrich was run with the flag "-a 35" and GEM run with the flags "-Xmx64G --k_min 8 --k_max 12." Once called, peaks were merged using HOMERs mergePeaks command with the flag "-d 500." Fold change over input of each peak at the weighted peak center was calculated using a custom python script with only uniquely mapped reads. Peaks were also annotated using HOMER's annotatePeaks command using the hg38 genome. For peak filtering to select for peaks within −2kb to +1kb of the TSS, the reported "Distance to TSS" value was used. Genome coverage plots were generated using ngsplot [17] with the flags "-G hg38 -R tss -L 2000 -RB 0.05" and tailored y-axis values per plot. To assess ChIP-Seq enrichment, a fingerprintPlot was generated using default parameters with deepTools.

To determine which putative SPEARs showed noticeable signal in their promoter, bedgraph counts were generated for gene promoters using HOMERs annotatePeaks command with the flags “tss hg38 -size 4000 -hist 25 -ghist.” These summed promoter intensities were then merged with the list of putative *SPEARs* to give a final list of confirmed *SPEARs* (External Database S1).

For generating the DMSO/DRB/ActD and siMYC coverage regions, bam coverage maps were generated using bamCoverage with the flags “--outFileFormat bedgraph --normalizeUsing None --binSize 25 --effectiveGenomeSize 2864785220.” The generated bedgraphs were then imported into R v4.0.3 (https://www.R-project.org/ and figures generated using the package Sushi ( https://github.com/dphansti/Sushi).

**Motif Discovery**

RNA binding motifs were identified by the following steps:

1) Filtering for *SPEARs* expression
2) Prediction of 5’ and 3’ *SPEARs* boundaries from RNA-Seq
3) Search for a common motif in selected *SPEARs*

In total, 13891 predicted promoter loci in HL60 (Broad ChromHMM) were subjected to coverage calculation. *SPEARs* were further selected according to their expression and the upper quartile subset ($75^{th}$ and above percentile) was chosen for further analysis. 5’ and 3’ *SPEARs* boundaries were inferred from RNA-Seq: coverage tracks were scanned and 5’ and 3’ *SPEARs* boundaries identified as a local drop in the level of coverage using Friedman’s SuperSmoother method (R, supsmu [21]) (Supplementary Data #4). In total, 2363 *SPEARs* were scanned for common motifs using the findGenomeMotif in RNA mode (Homer suite: http://homer.ucsd.edu/homer/ngs) with option “-len 10,20,30” using Human promoters as a background (except those scanned for motifs). Motifs were filtered according to both significance ($p<=e^{-10}$) and fold enrichment (observed vs expected >=200), leading to the identification of 3 enriched motifs (RM 3,5 and 9,).

The presence of the identified motifs in acH2A.Z and TIP60 RIP-Seq overlapping peaks was assessed by running findGenomeMotif with the –mknown option using non-overlapping peaks as background. The same analysis was carried out for SINE, LINE and rRNA transcripts. Repetitive elements were retrieved from UCSC (RepeatMasker) and transcripts expressed >= 2 fpkm were selected for motif scanning. rRNA genomic region were retrieved from the UCSC Table Browser (Table:rmsk, repClass:rRNA).

**Statistical Analysis:**

All the statistical analyses were performed using the R suite (https://www.r-project.org/). The statistical comparison of the distributions corresponding to ChIP-Seq, RIP-Seq and RNA-Seq cumulative read count signals were performed using the Mann Whitney Wilcoxon signed rank test, with a confidence interval of ($\wp < 0.05$).

**Data Availability**

Sequencing Data are available on the gene omnibus database under the accession ID number: GSE165526; enter token: slehymyczncvfqd. List of all data sets is in Supplementary Data #6. Mass Spectrometry data will be deposited to the ProteomeXChange Consortium via PRIDE upon acceptance of the manuscript. Private partial submission with MASSIVE: https://massive.ucsd.edu/ProteoSAFe/dataset.jsp?task=38a5eae58dd247f5b3c42f2be181e692; Password: Alex_Acetyl

**Supplementary Table A:** siRNA Sequences

| ***c-MYC SPEARs* siRNAs sequences** (sense strand): |
|---|
| 5'-GCGGAGGGAAAGACGCUUU-3' |
| 5'-CCAUCUUGAACAGCGUACA-3' |
| 5'-CGGCAAAGGCCUGGAGGCA-3' |
| 5'-GGGUAAUAACCCAUCUUGA-3' |
| 5'-GCUGAAUUGUGCAGUGCAU-3' |
| 5'-GGAUUUGGAAGCUACUAUA-3' |
| 5'-GGAAACCUUGCACCUCGGA-3' |
| 5'-GACAUCCAGGCGCGAUGAU-3' |
| 5'-CCAUUACCGGUUCUCCAUA-3' |
| 5'-GAAGCUACUAUAUUCACUU-3' |
| 5'-GAAAGACGCUUUGCAGCAA-3' |
| 5'-GGCCGUUUUAGGGUUUGUU-3' |
| 5'-GGCACACUUACUUUACUUU-3' |
| 5'-CGCUGAGCUGCAAACUCAA-3' |
| 5'-CCAACCUGAAAGAAUAACA-3' |
| 5'-GCGAUGAUCUCUGCUGCCA-3' |
| 5'-GUAAUUUGCAAUCCUUAAA-3' |
| 5'-GAGUAAUUUGCAAUCCUUA-3' |
| 5'-GUCUAUGUACUUGUGAAUU-3' |
| 5'-GCAAACUCAACGGGUAAUA-3' |
| 5'-GCAAAAUCCAGCAUAGCGA-3' |
| 5'-CAGUGCAUCGGAUUUGGAA-3' |
| 5'-GCAAUCCUUAAAGCUGAAU-3' |
| 5'-GGCUGGAAACUUGUUUUAA-3' |
| 5'-CUAUGUACUUGUGAAUUAU-3' |
| 5'-GCGUUUGCGGCAAAGGCCU-3' |
| 5'-GAACAGCGUACAUGCUAUA-3' |
| **Negative Control siPool (siControl**) |
| Sequences (sense strand): |
| 5'-UGUACGCGUCUCGCGAUUU-3' |
| 5'-UAUACGCGGUACGAUCGUU-3' |
| 5'-UUCGCGUAAUAGCGAUCGU-3' |
| 5'-UCGGCGUAGUUUCGACGAU-3' |
| 5'-UCGCGUAAGGUUCGCGUAU-3' |
| 5'-UCGCGAUUUUAGCGCGUAU-3' |
| 5'-UCGCGUAUAUACGCUACGU-3' |
| 5'-UUUCGCGAACGCGCGUAAU-3' |

| |
|---|
| 5'-UCGUAUCGUAUCGUACCGU-3' |
| 5'-UUAUCGCGCGUUAUCGCGU-3' |
| 5'-UCUCGUAGGUACGCGAUCU-3' |
| 5'-UCGUACUCGAUAGCGCAAU-3' |
| 5'-UUUGCGAUACCGUAACGCU-3' |
| 5'-UGCGUAAGGCAUGUCGUAU-3' |
| 5'-UUAUCGGCAGUUCGCCGUU-3' |
| 5'-UAGCGCGACAUCUAUCGCU-3' |
| 5'-UCGUCGUAUCAGCGCGUUU-3' |
| 5'-UACGCGAAACUGCGUUCGU-3' |
| 5'-UCGACGAUAGCUAUCGCGU-3' |
| 5'-UCGCGUAAUACGCGAUCGU-3' |
| 5'-UCGCGAUAAUGUUACGCGU-3' |
| 5'-UUAACGCGCUACGCGUAUU-3' |
| 5'-UCGCGUAUAGGUAACGCGU-3' |
| 5'-UUACGCGAUCACGUAACGU-3' |
| 5'-UUAUCGCGCGUCGCGUAAU-3' |
| 5'-UUACGUACUAGUGCGUACU-3' |
| 5'-UAUACGCCGGUUGCGUAGU-3' |
| 5'-UUCGCGUGCAUAGCGUAAU-3' |
| 5'-UACGCGACCUAAUCGCGAU-3' |
| 5'-UCGUACGCUGAACGCGUAU-3' |

**Supplementary Table B.** Primer-Sequences used for Chromatin Immunoprecipitation qPCR for *c-MYC* and *PU.1* genes

| | |
|---|---|
| Forward mychip1 | 5'-GGC TAA TCC TCT ATG GGA GTC TGT C-3' |
| Reverse mychip2 | 5'- TTT CTG AAT ACT AGT GAA AGT GCA-3' |
| Forward mychip3 | 5'- TCA GAA AAA ATT GTG AGT CAG TGA -3' |
| Reverse mychip4 | 5'- TTG TGG ACC GAG CCG GGG GAG TCA -3' |
| Forward mychip5 | 5'- CCG GCT CGG TCC ACA AGC TCT CCA -3' |
| Reverse mychip6 | 5'- TCT GCC TGT TCC AGA GCT GGG CTA -3' |
| Forward mychip7 | 5'- ACA GGC AGA CAC ATC TCA GGG CTA -3' |
| Reverse mychip8 | 5'- ATA GGG AGG AAT GAT AGA GGC ATA -3' |
| Forward mychip9 | 5'- CTA CAC TAA CAT CCC ACG CTC TGA -3' |
| Reverse mychip10 | 5'- AAC CGC ATC CTT GTC CTG TGA GTA -3' |
| Forward mychip11 | 5'- AAG GAT GCG GTT TGT CAA ACA GTA -3' |
| Reverse mychip12 | 5'- TCC TCA GCC GTC CAG ACC CTC GCA -3' |
| Forward mychip13 | 5'- TAG AGT GCT CGG CTG CCC GGC TGA -3' |
| Reverse mychip14 | 5'- TCT GAG AAG CCC TGC CCT TCT CGA -3' |
| Forward mychip15 | 5'- GAA CGG AGG GAG GGA TCG CGC TGA -3' |
| Reverse mychip16 | 5'- GTG CAA AGT GCC CGC CCG CTG CTA -3' |

| | |
|---|---|
| Forward mychip17 | 5'- GAC TCT CCC GAC GCG GGG AGG CTA -3' |
| Reverse mychip18 | 5'- CCC CAG TTA CCA TAA CTA CTC TGA -3' |
| Forward mychip19 | 5'- GGA TCG GGG TAA AGT GAC TTG TCA -3' |
| Reverse mychip20 | 5'- GCG GCT GCG GAG CGA TCT GGC TCA -3' |
| Forward mychip21 | 5'- GCC AGA TCG CTC CGC AGC CGC TGA -3' |
| Reverse mychip22 | 5'- ACA CCA CGT CCT AAC ACC TCT AGA -3' |
| Forward PU.1ChIP | 5'- AAA GTC ATC CCT CTC AGT CCC AGC -3' |
| Reverse PU.1 ChIP | 5'- GAA GGG CCT GCC GCT GGG AGA TAG -3' |

**Supplementary Table C.** RNA/DNA Oligonucleotides and peptides used for REMSA:

| | |
|---|---|
| RM9A | 5’-GGC GUG GCG GUG GGC GCG CAG U-3’ |
| MutRM9A | 5’-UUA UGU UAU UGU UUA UAU ACG U-3’ |
| Unrelated (UR) | 5’-GCG CCC UGC AGC CUG GUA CGC G-3’ |
| Unrelated 2 (UR2) | 5’-CUU UCC UCC ACU CUC CCU GGG A-3’ |
| Unrelated 3 (UR3) | 5’-GCC CUU UCC CCA GCC UUA GCG A-3’ |
| Mut1_UR2 | 5’-CUU UCA GAA CAG AGA CCU GGG A-3’ |
| Mut2_UR2 | 5’-CUU UCU CUU GUC UCU CCU GGG A-3’ |
| Mut_UR3 | 5’-GCC CUU UAA AAC UAA GGA GCG A-3’ |
| DM9F | 5’-GCC CTT TCC CCA GCC TTA GCG A-3’ |
| DM9R | 5’-TCG CTA AGG CTG GGG AAA GGG C-3’ |
| DM2F | 5’-CTT TCC TCC ACT CTC CCT GGG A-3’ |
| DM2R | 5’-TCC CAG GGA GAG TGG AGG AAA G-3’ |
| DM3F | 5’-GGC GTG GCG GTG GGC GCG CAG T-3’ |
| DM3R | 5’-ACT GCG CGC CCA CCG CCA CGC C-3’ |
| H2A.Z peptide | H-AGGKAGKDSGKAKTKAVSRS-OH |
