## Supplementary material for "Formation and recycling of an active epigenetic mark mediated by cell cycle-specific RNAs": Supl.5

| Protein FDR Confid | Master | Accession |
|---|---|---|
| High | Master Protein | P38507 |
| High | Master Protein | P00766 |
| High | Master Protein | P02769 |
| High | Master Protein | P35527 |
| High | Master Protein | O46375 |
| High | Master Protein | P04264 |
| High | Master Protein | P01870 |
| High | Master Protein | 478694 |
| High | Master Protein | P12763 |
| High | Master Protein | P0C0S5 |
| High | Master Protein | 88041 |
| High | Master Protein | P02666 |
| High | Master Protein | P15636 |
| Medium | Master Protein | P15497 |
| High | Master Protein | P02768 |
| Medium | Master Protein | P06396 |
| Medium | Master Protein | P35908 |
| Medium | Master Protein | P22629 |
| Medium | Master Protein | P02663 |
| Medium | Master Protein | P81605 |
| Low | Master Protein | Q86YZ3 |
| Medium | Master Protein | Q2UVX4 |
| High | Master Protein | 106849 |
| Medium | Master Protein | Q5D862 |
| Low | Master Protein | Q29443 |
| Medium | Master Protein | 1708589 |
| Medium | Master Protein | Q15149 |
| Medium | Master Protein | 1346345 |
| Medium | Master Protein | P01696 |
|  | Master Protein | P05109 |
| Medium | Master Protein | Q9NSB2 |
| Medium | Master Protein | 547753 |
|  | Master Protein | Q59146 |
|  | Master Protein | Q8N9T8 |
| Medium | Master Protein | 547751 |
|  | Master Protein | P15252 |
| Medium | Master Protein | P08670 |
| Low | Master Protein | P00761 |
|  | Master Protein | Q29461 |
|  | Master Protein | P08659 |
|  | Master Protein | P23254 |
| Medium | Master Protein | Q9Y446 |
| Medium | Master Protein | 125108 |
| Medium | Master Protein | 125098 |
|  | Master Protein | 37812657 |
| High | Master Protein | P13645 |
| Medium | Master Protein | 547752 |
| Medium | Master Protein | P02533 |
| Medium | Master Protein | Q6A163 |
| Low | Master Protein | Q92817 |
| Low | Master Protein | Q86SJ6 |
| Low | Master Protein | O60437 |
| Low | Master Protein | P32926 |
| Low | Master Protein | 327282624 |

| Description | Contamina |
|---|---|
| *CON* Immunoglobulin G-binding protein A OS=Staphylococcus aureus GN=spa PE=1 SV=1 | TRUE |
| *CON* Chymotrypsinogen A - Bos taurus | TRUE |
| *CON* serum albumin precursor [Bos taurus] | TRUE |
| *CON* Keratin, type I cytoskeletal 9 OS=Homo sapiens GN=KRT9 PE=1 SV=3 | TRUE |
| *CON* Transthyretin OS=Bos taurus GN=TTR PE=2 SV=1 | TRUE |
| *CON* Keratin, type II cytoskeletal 1 OS=Homo sapiens GN=KRT1 PE=1 SV=6 | TRUE |
| *CON* Ig gamma chain C region OS=Oryctolagus cuniculus PE=1 SV=1 | TRUE |
| S25705 Ig mu chain - sheep | TRUE |
| *CON* Alpha-2-HS-glycoprotein precursor - Bos taurus | TRUE |
| Histone H2A.Z OS=Homo sapiens OX=9606 GN=H2AFZ PE=1 SV=2 | FALSE |
| A31994 *CON* keratin 10, type I, epidermal - human gi623409 (J04029) keratin 10 [Homo sapiens] | TRUE |
| *CON* Beta-casein precursor - Bos taurus | TRUE |
| *CON* PROTEASE I PRECURSOR (API) (LYSYL ENDOPEPTIDASE). | TRUE |
| *CON* Apolipoprotein A-I OS=Bos taurus GN=APOA1 PE=1 SV=3 | TRUE |
| *CON* serum albumin precursor [Homo sapiens]. | TRUE |
| *CON* Gelsolin OS=Homo sapiens GN=GSN PE=1 SV=1 | TRUE |
| *CON* Keratin, type II cytoskeletal 2 epidermal OS=Homo sapiens GN=KRT2 PE=1 SV=2 | TRUE |
| *CON* Streptavidin OS=Streptomyces avidinii PE=1 SV=1 | TRUE |
| *CON* Alpha-S2-casein precursor - Bos taurus | TRUE |
| *CON* Dermcidin precursor - Homo sapiens | TRUE |
| *CON* Hornerin - Homo sapiens | TRUE |
| *CON* Complement C3 OS=Bos taurus GN=C3 PE=1 SV=2 | TRUE |
| PC1102 *CON* keratin 10, type I, cytoskeletal (clone HK51) - human (fragment) gi186629 (M77663) k | TRUE |
| *CON* Filaggrin-2 OS=Homo sapiens GN=FLG2 PE=1 SV=1 | TRUE |
| *CON* Serotransferrin OS=Bos taurus GN=TF PE=2 SV=1 | TRUE |
| *CON* Keratin, type i cytoskeletal 16 (cytokeratin 16) (K16) (CK 16) | TRUE |
| *CON* Plectin-1 OS=Homo sapiens GN=PLEC1 PE=1 SV=3 | TRUE |
| *CON* Keratin, type ii cytoskeletal 6b (cytokeratin 6b) (CK 6b) (k6b keratin) gi2119220pirI61767 kerati | TRUE |
| *CON* Ig kappa chain V region K29-213 OS=Oryctolagus cuniculus PE=1 SV=1 | TRUE |
| *CON* Protein S100-A8 OS=Homo sapiens GN=S100A8 PE=1 SV=1 | TRUE |
| *CON* Keratin, type II cuticular Hb4 OS=Homo sapiens GN=KRT84 PE=1 SV=2 | TRUE |
| *CON* Keratin, type ii cytoskeletal 4 (cytokeratin 4) (K4) (CK4) | TRUE |
| *CON* Glucan endo-1,3-beta-glucosidase (Zymolase) OS=Arthrobacter sp. (strain YCWD3) GN=glcI I | TRUE |
| *CON* Protein KRI1 homolog OS=Homo sapiens GN=KRI1 PE=1 SV=2 | TRUE |
| *CON* Keratin, type i cytoskeletal 17 (cytokeratin 17) (K17) (CK 17) (39.1) (VERSION 1) gi422802pirS | TRUE |
| *CON* Rubber elongation factor protein OS=Hevea brasiliensis PE=1 SV=2 | TRUE |
| *CON* Vimentin OS=Homo sapiens GN=VIM PE=1 SV=4 | TRUE |
| *CON* Trypsin precursor - sus scrofa - includes MSPRL added real and artificial sequences to mimic | TRUE |
| *CON* Elastase-2A precursor - Bos taurus | TRUE |
| *CON* Luciferin 4-monooxygenase Firefly luciferase OS=Photinus pyralis PE=1 SV=1 | TRUE |
| *CON* Transketolase 1 - Saccharomyces cerevisiae | TRUE |
| *CON* Plakophilin-3 OS=Homo sapiens GN=PKP3 PE=1 SV=1 | TRUE |
| *CON* Keratin, type ii cytoskeletal 7 (cytokeratin 7) (K7) (CK 7) gi1200072 (X13320) keratin [Homo sa | TRUE |
| *CON* Keratin, type ii cytoskeletal 3 (cytokeratin 3) (K3) (CK3) (65 KD cytokeratin) | TRUE |
| beta galactosidase [UAS-less reporter vector pMELbeta] gi\|37812664\|gb\|AAR04150.1\| beta galactosic | TRUE |
| *CON* Keratin, type I cytoskeletal 10 OS=Homo sapiens GN=KRT10 PE=1 SV=4 | TRUE |
| *CON* Keratin, type ii cytoskeletal 2 oral (cytokeratin 2P) (K2P) (CK 2P) gi2119218pirI53169 cytokera | TRUE |
| *CON* Keratin, type I cytoskeletal 14 OS=Homo sapiens GN=KRT14 PE=1 SV=3 | TRUE |
| *CON* Keratin, type I cytoskeletal 39 OS=Homo sapiens GN=KRT39 PE=1 SV=2 | TRUE |
| *CON* Envoplakin OS=Homo sapiens GN=EVPL PE=1 SV=2 | TRUE |
| *CON* Desmoglein-4 OS=Homo sapiens GN=DSG4 PE=1 SV=1 | TRUE |
| *CON* Periplakin OS=Homo sapiens GN=PPL PE=1 SV=2 | TRUE |
| *CON* Desmoglein-3 OS=Homo sapiens GN=DSG3 PE=1 SV=2 | TRUE |
| PREDICTED: translational activator GCN1-like [Anolis carolinensis] | TRUE |

| Exp. q-valu | Exp. q-valu | Coverage [ | # Peptides | # PSMs | # Unique P | # AAs | MW [kDa] | calc. pI | Score Sequ |
|---|---|---|---|---|---|---|---|---|---|
| | 0 | 74 | 348 | 1891 | 348 | 508 | 55.4 | 5.71 | 3226.77 |
| | 0 | 66 | 60 | 249 | 60 | 245 | 25.7 | 8.16 | 356.56 |
| | 0 | 47 | 53 | 126 | 49 | 607 | 69.2 | 6.11 | 160.53 |
| | 0 | 24 | 18 | 37 | 18 | 623 | 62 | 5.24 | 57.37 |
| | 0 | 70 | 17 | 55 | 17 | 147 | 15.7 | 6.3 | 81.65 |
| | 0 | 24 | 16 | 37 | 15 | 644 | 66 | 8.12 | 58 |
| | 0 | 33 | 16 | 69 | 16 | 323 | 35.4 | 8.29 | 42.3 |
| | 0 | 18 | 10 | 28 | 10 | 592 | 64.5 | 5.71 | 32.53 |
| | 0 | 33 | 10 | 16 | 10 | 359 | 38.4 | 5.5 | 21.77 |
| | 0 | 60 | 10 | 18 | 10 | 128 | 13.5 | 10.58 | 29.62 |
| | 0 | 25 | 10 | 29 | 1 | 561 | 57.2 | 5.07 | 38.62 |
| | 0 | 40 | 7 | 11 | 7 | 224 | 25.1 | 5.35 | 14.33 |
| | 0 | 10 | 6 | 13 | 6 | 653 | 68.1 | 7.24 | 18.84 |
| | 0.033 | 15 | 3 | 9 | 3 | 265 | 30.3 | 5.97 | 11.37 |
| | 0 | 14 | 11 | 22 | 7 | 609 | 69.3 | 6.28 | 48.11 |
| | 0.029 | 4 | 3 | 7 | 3 | 782 | 85.6 | 6.28 | 8.47 |
| | 0.021 | 10 | 3 | 4 | 2 | 639 | 65.4 | 8 | 6.26 |
| | 0.022 | 11 | 2 | 5 | 2 | 183 | 18.8 | 8.35 | 6.28 |
| | 0.012 | 9 | 2 | 3 | 2 | 222 | 26 | 8.43 | 3.92 |
| | 0.024 | 26 | 2 | 4 | 2 | 110 | 11.3 | 6.54 | 3.29 |
| | 0.063 | 7 | 4 | 6 | 3 | 2850 | 282.2 | 10.04 | 2.26 |
| | 0.031 | 3 | 4 | 6 | 4 | 1661 | 187.1 | 6.84 | 9.28 |
| | 0 | 26 | 9 | 15 | 1 | 384 | 39.7 | 4.78 | 35.63 |
| | 0.027 | 4 | 3 | 5 | 2 | 2391 | 247.9 | 8.31 | 7.9 |
| | 0.059 | 2 | 1 | 2 | 1 | 704 | 77.7 | 7.08 | 2.07 |
| | 0.024 | 5 | 2 | 2 | 1 | 473 | 50.9 | 5.44 | 3.21 |
| | 0.012 | 1 | 3 | 3 | 3 | 4684 | 531.5 | 5.96 | 4.24 |
| | 0.024 | 10 | 4 | 5 | 2 | 564 | 60 | 8 | 7.32 |
| | 0.021 | 9 | 1 | 2 | 1 | 110 | 11.6 | 8.97 | 3.02 |
| | | 12 | 1 | 1 | 1 | 93 | 10.8 | 7.03 | |
| | 0.015 | 8 | 3 | 3 | 1 | 600 | 64.8 | 7.56 | 5.06 |
| | 0.029 | 4 | 2 | 3 | 1 | 534 | 57.2 | 6.61 | 8.43 |
| | | 6 | 1 | 1 | 1 | 548 | 58.1 | 7.43 | |
| | | 4 | 1 | 1 | 1 | 709 | 83.2 | 5.17 | |
| | 0.02 | 6 | 2 | 3 | 1 | 432 | 48.1 | 5.02 | 6.04 |
| | | 13 | 1 | 1 | 1 | 80 | 8.7 | 4.59 | |
| | 0.02 | 5 | 2 | 2 | 1 | 466 | 53.6 | 5.12 | 2.83 |
| | 0.057 | 4 | 1 | 2 | 1 | 741 | 79.4 | 8.31 | 1.99 |
| | | 4 | 1 | 1 | 1 | 269 | 28.8 | 8.34 | |
| | | 3 | 1 | 1 | 1 | 550 | 60.7 | 6.9 | |
| | | 2 | 1 | 1 | 1 | 680 | 73.8 | 7.01 | |
| | 0.016 | 3 | 2 | 3 | 2 | 797 | 87 | 9.32 | 5.06 |
| | 0.013 | 12 | 3 | 3 | 1 | 469 | 51.3 | 5.49 | 5.06 |
| | 0.013 | 8 | 3 | 3 | 1 | 629 | 64.5 | 6.48 | 5.06 |
| | | 3 | 1 | 1 | 1 | 1045 | 118.5 | 5.8 | |
| | 0 | 21 | 12 | 18 | 2 | 593 | 59.5 | 5.21 | 43.72 |
| | 0.03 | 6 | 3 | 3 | 1 | 638 | 65.8 | 8.12 | 9.15 |
| | 0.019 | 6 | 2 | 2 | 1 | 472 | 51.6 | 5.16 | 6.03 |
| | 0.021 | 5 | 1 | 1 | 1 | 491 | 55.6 | 5.26 | 2.87 |
| | 0.066 | 1 | 2 | 2 | 2 | 2033 | 231.5 | 6.96 | 2.56 |
| | 0.065 | 3 | 1 | 1 | 1 | 1040 | 113.8 | 4.56 | 2.34 |
| | 0.062 | 1 | 1 | 1 | 1 | 1756 | 204.5 | 5.57 | 2.25 |
| | 0.058 | 1 | 1 | 1 | 1 | 999 | 107.5 | 5 | 2.06 |
| | 0.137 | 0 | 1 | 1 | 1 | 2671 | 293.5 | 7.27 | 1.78 |

| # Peptides | # Peptides | Found in S | Found in S | \|Log Prob\| | # Protein G | # Cumulati | FDR (by Se | Confidence (by Search |
|---|---|---|---|---|---|---|---|---|
| 339 | 305 | High | High | 2735.04 | 1 | 0 | 0 | 3 |
| 55 | 52 | Not Found | High | 316.84 | 1 | 0 | 0 | 3 |
| 49 | 46 | High | High | 219.81 | 1 | 0 | 0 | 3 |
| 16 | 16 | High | High | 105.4 | 1 | 0 | 0 | 3 |
| 16 | 16 | High | High | 104.58 | 1 | 0 | 0 | 3 |
| 14 | 15 | High | High | 70.38 | 1 | 0 | 0 | 3 |
| 15 | 13 | Not Found | High | 65.9 | 1 | 0 | 0 | 3 |
| 10 | 7 | High | High | 45.06 | 1 | 0 | 0 | 3 |
| 7 | 6 | High | High | 42.66 | 1 | 0 | 0 | 3 |
| 8 | 7 | High | High | 38.36 | 1 | 0 | 0 | 3 |
| 8 | 9 | High | High | 37.8 | 1 | 0 | 0 | 3 |
| 6 | 5 | High | High | 36.95 | 1 | 0 | 0 | 3 |
| 6 | 5 | High | High | 31.91 | 1 | 0 | 0 | 3 |
| 3 | 3 | High | Not Found | 21.12 | 1 | 0 | 0 | 3 |
| 4 | 11 | High | High | 18.46 | 1 | 0 | 0 | 3 |
| 3 | 3 | Not Found | High | 17.25 | 1 | 0 | 0 | 3 |
| 2 | 2 | High | High | 10 | 1 | 0 | 0 | 3 |
| 2 | 2 | Not Found | High | 9.64 | 1 | 0 | 0 | 3 |
| 2 | 1 | High | High | 9.49 | 1 | 0 | 0 | 3 |
| 2 | 1 | High | Not Found | 8.61 | 1 | 0 | 0 | 3 |
| 3 | 2 | Not Found | High | 8.5 | 1 | 0 | 0 | 3 |
| 2 | 4 | Not Found | High | 6.99 | 1 | 0 | 0 | 3 |
| 1 | 9 | Not Found | High | 6.21 | 1 | 0 | 0 | 3 |
| 2 | 3 | High | High | 5.21 | 1 | 0 | 0 | 3 |
| 1 | 1 | Not Found | High | 5 | 1 | 1 | 0.04 | 2 |
| 1 | 1 | Not Found | High | 4.88 | 1 | 1 | 0.038462 | 2 |
| 1 | 2 | High | High | 4.73 | 1 | 1 | 0.037037 | 2 |
| 2 | 3 | High | Not Found | 4.51 | 1 | 1 | 0.035714 | 2 |
| 1 | 1 | Not Found | High | 4.3 | 1 | 1 | 0.034483 | 2 |
| 1 |  | High | Not Found | 3.71 | 1 | 1 | 0.033333 | 2 |
| 1 | 2 | High | Not Found | 3.7 | 1 | 1 | 0.032258 | 2 |
|  | 2 | Not Found | High | 3.64 | 1 | 1 | 0.03125 | 2 |
| 1 |  | High | Not Found | 3.22 | 1 | 2 | 0.060606 | 1 |
| 1 |  | High | Not Found | 3.06 | 1 | 3 | 0.088235 | 1 |
| 1 | 2 | High | Not Found | 2.89 | 1 | 3 | 0.085714 | 1 |
| 1 |  | High | Not Found | 2.72 | 1 | 3 | 0.083333 | 1 |
| 1 | 1 | High | Not Found | 2.54 | 1 | 3 | 0.081081 | 1 |
| 1 | 1 | Not Found | High | 2.26 | 1 | 4 | 0.105263 | 1 |
| 1 |  | High | Not Found | 2.14 | 1 | 4 | 0.102564 | 1 |
| 1 |  | Not Found | High | 1.7 | 1 | 8 | 0.195122 | 1 |
| 1 |  | High | Not Found | 1.64 | 1 | 8 | 0.190476 | 1 |
| 1 | 2 | High | High | 1.62 | 1 | 8 | 0.186047 | 1 |
| 1 | 2 | Not Found | High | 1.16 | 1 | 9 | 0.204545 | 1 |
| 1 | 2 | High | Not Found | 0.95 | 1 | 10 | 0.222222 | 1 |
| 1 |  | Not Found | High | 0.7 | 1 | 12 | 0.25 | 1 |
|  | 12 | High | High |  | 1 | 0 | 0 | 0 |
|  | 3 | Not Found | High |  | 1 | 0 | 0 | 0 |
|  | 2 | Not Found | High |  | 1 | 0 | 0 | 0 |
|  | 1 | High | Not Found |  | 1 | 0 | 0 | 0 |
|  | 2 | Not Found | High |  | 1 | 0 | 0 | 0 |
|  | 1 | High | Not Found |  | 1 | 0 | 0 | 0 |
|  | 1 | High | Not Found |  | 1 | 0 | 0 | 0 |
|  | 1 | Not Found | High |  | 1 | 0 | 0 | 0 |
|  | 1 | Not Found | High |  | 1 | 0 | 0 | 0 |

ı Engine): PMI-Byonic A6
